# Termination sequence between an inducible promoter and ubiquitous chromatin opening element (UCOE) reduces gene expression leakage and silencing

**DOI:** 10.1101/2024.12.18.629244

**Authors:** Tomoki Yanagi, Shean Fu Phen, Jonah Ayala, Deniz Ece Aydin, David M. Truong

## Abstract

Inducible gene expression circuits offer precise control over target gene activation, making them essential tools for direct reprogramming, where cells are guided to differentiate into specific cell types. However, stable circuit function and consistent expression of key inducible proteins are crucial for effective reprogramming, and DNA methylation-induced silencing hinders this stability. To address this issue, A2-ubiquitous chromatin opening elements (A2UCOE) have gained attention for their ability to maintain open chromatin and prevent methylation, thereby stabilizing gene expression. In this study, we aimed to forward program iPSCs into thymic epithelial cells (TECs) using a compact, all-in-one gene circuit composed of a doxycycline-inducible Tet-On system, 863-bp A2UCOE (0.9 UCOE), and *FOXN1*, a master transcription factor for TEC differentiation. This compact construct enables site-specific genome integration and stable inducible expression within iPSCs for renewably generating TECs. While the 0.9 UCOE promoted stable expression of constitutively expressed genes, it also caused unintended *FOXN1* gene leakage, leading to unprogrammed gradual differentiation of the iPSCs. We generated A2UCOE fragments of varying lengths and found that gene leakage persisted regardless of fragment size. We tested spacer sequences between the A2UCOE and the Tet-On promoter consisting of varying AT-nucleotide content (35, 50, and 65%) and the 65% AT-rich SV40 poly-A terminator sequence and found that only the SV40 poly-A mitigated this leakage, and surprisingly, enhanced desired anti-silencing effects. These findings highlight the benefits and potential risks of using A2UCOE in iPSCs, and provides insights into a regulatory DNA architecture optimal for compact forward programming circuits for controlled iPSC differentiation.

**Graphical abstract:** 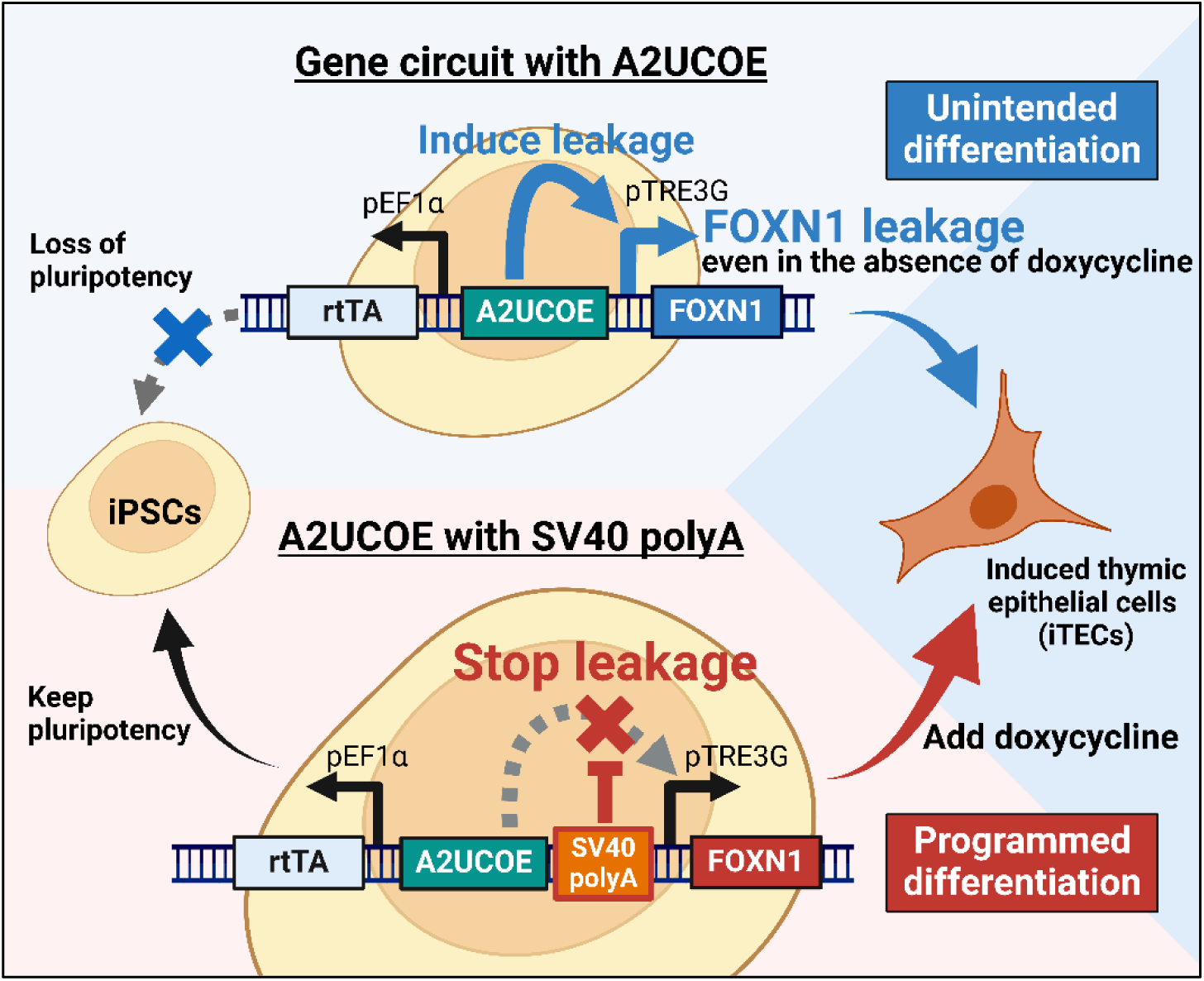

## Introduction

Mammalian inducible gene expression systems enable controlled expression of target genes in response to small, cell-permeable molecules such as doxycycline, and are widely utilized across many biological disciplines. One area where inducible systems are particularly essential is the field of direct reprogramming. Direct reprogramming or transdifferentiation is a rapidly emerging technology that enables the direct conversion of somatic cells into other lineages of cell types through the forced overexpression of master pioneer transcription factors (1). This technique has also been increasingly applied to hasten and aid the induction of renewable cell types like induced pluripotent stem cells (iPSCs) into desired somatic cells in what is called forward programming, since conventional directed differentiation approaches are lengthy and complex (2). Inducibility ensures that the iPSCs may be continuously cultured as a renewable resource, before committing to a somatic cell identity that has limited capacity for expansion.

The most widely used inducible system for direct reprogramming is the reverse tetracycline-controlled transactivator (rtTA) system, commonly known as the Tet-On system (3,4). In the Tet-On system, exogenously applied doxycycline, a more potent derivative of tetracycline, complexes to rtTA enabling it to bind and activate gene expression at Tet binding sites found in synthetic tetracycline response element (TRE) promoters. However, the Tet-On system is entirely dependent on stable expression of rtTA within cells (5,6), as changes in rtTA expression levels reduce or even eliminate output (7). Transgene silencing of rtTA, presumably through DNA methylation, has been a common problem, especially in iPSCs, which express high levels of de novo DNA methyltransferases (8,9).

Most direct reprogramming Tet-On gene circuits utilize a split design, where the rtTA components are expressed from a constitutive promoter integrated at one location, while the target genes under the rtTA-responsive promoters are integrated at another location (10). Genomic integrations are typically performed using lentivirus particles, which have a limited DNA size capacity (<10 kb) and integrates randomly in the genome (11,12). The random integration often results in position-effect variegation of expression and complete silencing over time (13,14). Site-specific integration into separate safe-harbor genomic sites like ROSA26 or AAVS1 via homology-directed repair help mitigate this in a clonal manner (15), but the clones largely still suffer the same eventual transgene silencing effects. Alternatively, "all-in-one" versions of the Tet-On system, which join the constitutive and inducible elements in a bidirectional transcriptionally opposing configuration, enable a more compact circuit that offers more stable expression (10).

Even so, these systems often include insulator elements to help mitigate silencing even further. The A2-ubiquitous chromatin opening element (A2UCOE) has been shown to reduce gene silencing when paired to diverse constitutive promoters and in different cell types (16–18). The A2UCOE element is located on human chromosome 7, between the housekeeping genes heterogeneous nuclear ribonucleoprotein A2/B1 (*HNRNAPA2B1*) and chromobox 3 (*CBX3*), which are transcribed in opposite directions. The A2UCOE region spans approximately 3 kb and contains a large CpG island, which is unusual in its density of CpG dinucleotides. While most CpG islands attract DNA methylation, the CpG density in the A2UCOE is thought to resist methylation by maintaining an open chromatin structure, thereby helping prevent gene silencing through epigenetic repression mechanisms (19–21). Further studies have revealed shorter lengths of the A2UCOE, from 2.2 kb down to even 0.6 kb, that also effectively reduce silencing (16,17,21–31). While it remains under investigation which length is most effective for stable gene expression, these shorter A2UCOE fragments have promise in compact gene circuits such as those used in gene therapy.

Here, we constructed an "all-in-one" Tet-On system utilizing the A2UCOE that is optimized for site-specific genome integration through homology-directed repair. As a model for studying the transcription dynamics of our system, we integrated and express the pioneer transcription factor *FOXN1* in human iPSCs. When constitutively overexpressed, *FOXN1* is a powerful pioneer factor that is sufficient to force fibroblasts and iPSCs into thymic epithelial cells (TECs), which make up the thymus where T-cells are matured (32–34). We hypothesized that an inducible *FOXN1* expression system would enable a renewable source of human TECs for pre-clinical studies. However, our research demonstrates that while the A2UCOE does reduce silencing of the constitutively expressed portions of the construct, it also causes leaky expression of the TRE promoter directed genes. To date, while A2UCOE has been recognized for its contribution to stabilizing gene expression, the potential risks associated with its use in gene expression have not been documented. We demonstrate that incorporating an SV40 polyadenylation (poly-A) spacer element between the UCOE and the inducible TRE promoter alleviates this issue and is not likely due to its AT nucleotide content. This study underscores both the benefits and possible limitations of using A2UCOE and introduces a novel design consideration for regulatory DNA arrangements in mammalian gene circuit design.

## Results

### Development of the "all-in-one" Tet-On system

We developed an "all-in-one" Tet-On system, a gene circuit containing a Tet-On control mechanism on a single plasmid, which facilitates straightforward knock-in of the gene circuit into target genomic regions. The design of the all-in-one Tet-On plasmid is shown in **Figure 1A**. Downstream of the human EF1α promoter, we placed the rtTA gene, which controls gene transcription by binding to the TRE3G promoter, along with a blasticidin resistance (BSR) gene for selection and the mScarlet (mScar) reporter gene. The TRE3G promoter is positioned in the opposite orientation to the EF1α promoter, enabling the insertion of a target gene downstream of the TRE3G promoter for regulated expression of a gene of interest. The TRE3G promoter allows for target gene expression in the presence of both rtTA and doxycycline.

**Figure 1.**
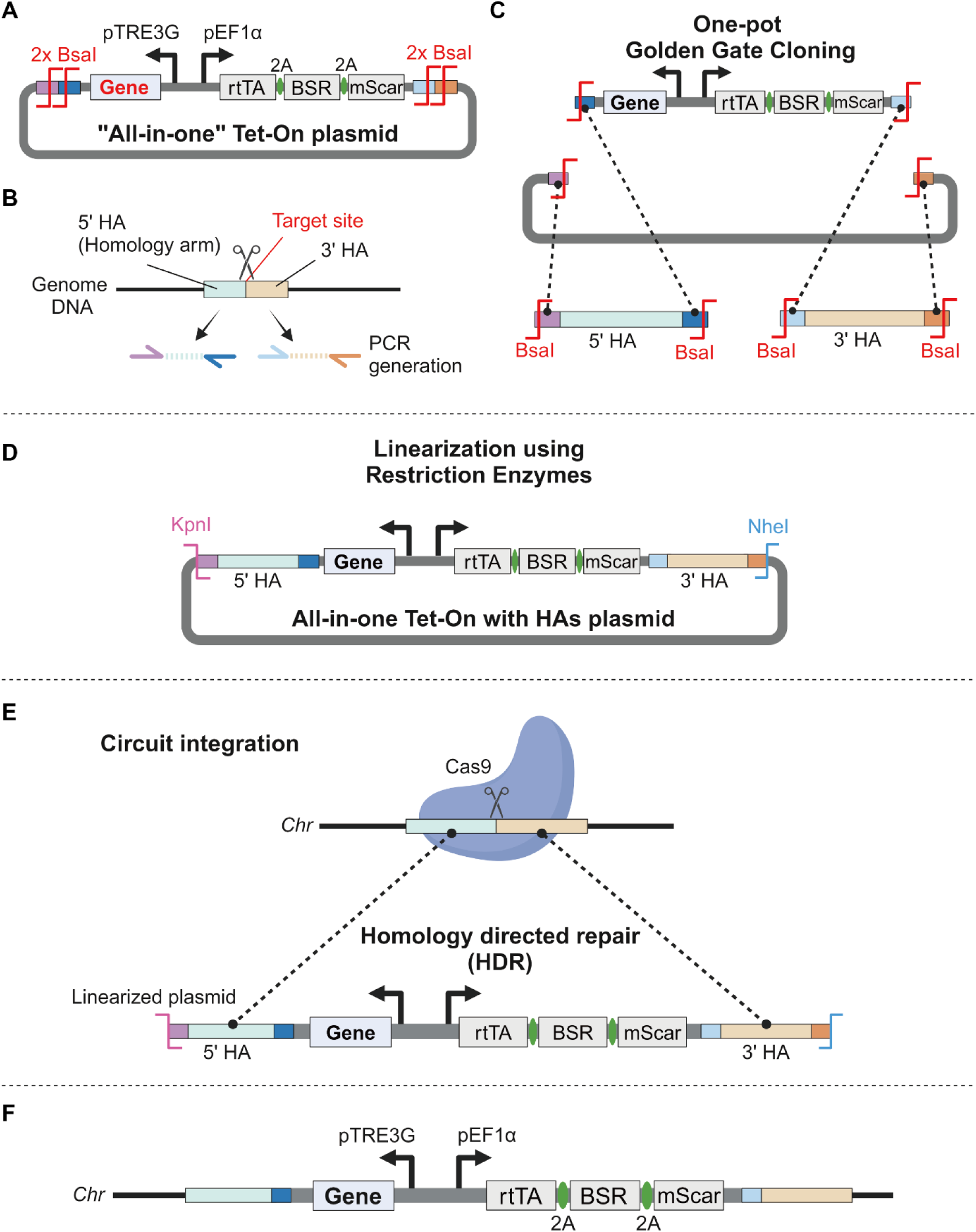
Design and integration of the "all-in-one" Tet-On system plasmid for targeted gene knock-in. A. Schematic representation of the "all-in-one" Tet-On plasmid. The human EF1α promoter drives the expression of the reverse tetracycline transactivator (rtTA), along with the blasticidin resistance (BSR) gene and mScarlet (mScar) reporter gene. The TRE3G promoter is oriented in the opposite direction to control the expression of the target gene (Gene). Two sets of adjacent BsaI restriction sites (2x BsaI) flank the regulatory elements. B. Generation of homology arms (HAs) through PCR amplification. The 5’ and 3’ homology arms, each approximately 450 bp, are designed to match the target genomic site for precise knock-in through homologous recombination. C. One-pot Golden Gate Cloning is used to insert the homology arms into the all-in-one plasmid. The BsaI restriction enzyme is used to ligate the homology arms with the Tet-On gene circuit. D. The plasmid (All-in-one Tet-On with HRs plasmid) is linearized using KpnI and NheI restriction enzymes to prepare it for integration into the genome. The final construct includes the homology arms and the regulatory elements of Tet-On system. E. Schematic of the targeted circuit integration. The linearized plasmid and a Cas9-gRNA vector are co-electroporated into mammalian cells. Cas9 creates a double-stranded break at the target site, and the homology arms direct homologous recombination (HDR), enabling precise knock-in of the gene circuit into the desired genomic location. F. Final structure of the genomic integration, with the All-in-one Tet-On gene circuit at the target site.

To facilitate precise genomic integration, we incorporated two pairs of adjacent BsaI restriction sites flanking the regulatory elements on the plasmid to allow for scarless Golden Gate cloning. As shown in **Figure 1B**, we first identified the cleavage site in the target genomic region using CRISPR/Cas9 and designed homology arms (HA) of approximately 450 bp for the 5’ and 3’ regions, which were then amplified by PCR. Next, we used the BsaI restriction enzyme and one-pot Golden Gate cloning to insert these amplified HAs into the all-in-one Tet-On plasmid (**Figure 1C**), resulting in a construct with HAs flanking the gene expression circuit. This plasmid design also allows for easy linearization using the KpnI and NheI restriction enzymes (**Figure 1D**).

The plasmid, linearized using KpnI and NheI, was then co-electroporated with a Cas9-gRNA-expressing plasmid into mammalian cells to induce homology-directed repair (HDR) (**Figure 1E**), thereby facilitating the targeted knock-in of the gene cassette into the desired genomic region (**Figure 1F**). This approach enabled us to develop an "all-in-one" plasmid capable of effectively controlling the expression of target genes in potentially any genomic region using a drug-inducible gene circuit.

### 0.9 UCOE enhances long-term gene expression in gene-edited iPS cells

The A2UCOE is located on human chromosome 7, between the *HNRNAPA2B1* and *CBX3* genes. This region is known as a CpG island, rich in CpG dinucleotides, and is generally resistant to methylation (19,20). We analyzed ATAC-seq data of this region in PGP1 iPSCs from the ENCODE consortium (35), and observed peaks corresponding to the CpG island, suggesting high transcriptional activity in the same region within PGP1 cells (**Figure 2A**). From this region, we isolated a UCOE fragment of 863 bp, which includes exon1 of *CBX3* and part of the *HNRNAPA2B1* promoter region, and termed it 0.9 UCOE (**Figure 2A**, **Supplemental Table 1**). Using this 0.9 UCOE, we constructed the all-in-one plasmid to induce the expression of codon-optimized human *FOXN1*, a key regulator of thymic epithelial cell (TEC) differentiation (**Figure 2B**) (32,36). TECs are critical players in both positive and negative selection of precursor T-cells during central tolerance and adaptive immunity (37). To minimize gene silencing and promote stable long-term expression of rtTA and the selection gene, we placed 0.9 UCOE upstream of the constitutive human EF1α promoter, which drives the expression of rtTA and selection genes, including BSR and mScarlet genes (**Figure 2B**). This gene circuit was integrated into the safe harbor region on chromosome 1 (Rogi1) of PGP1 human iPSCs using CRISPR/Cas9-assisted HDR (38,39). As a control, we created a gene circuit lacking UCOE (w/o UCOE) and integrated it into PGP1 cells similarly.

**Figure 2.**
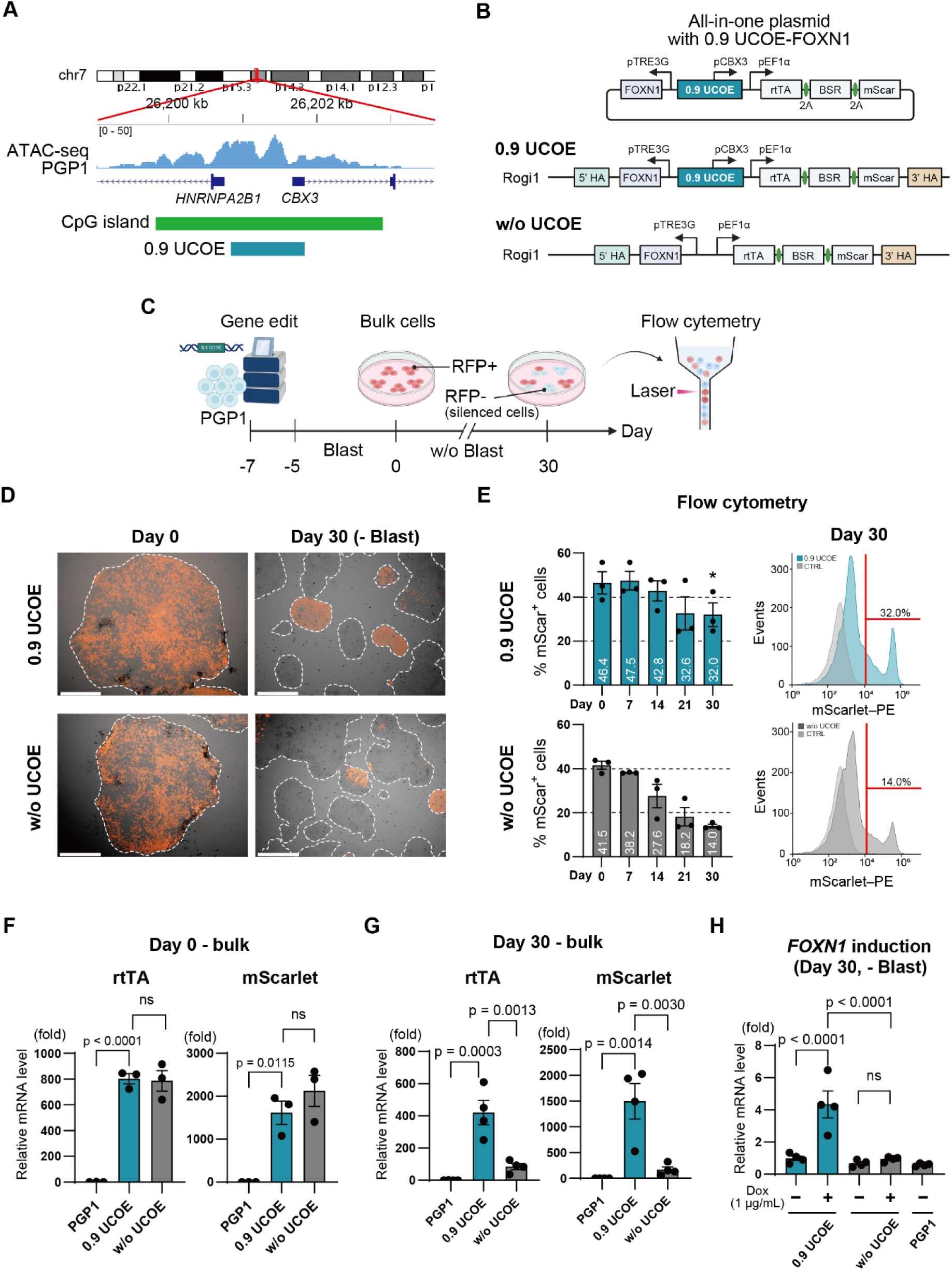
The 0.9 UCOE enhances long-term gene expression stability in gene-edited iPS cells. A. Location and design of the 0.9 UCOE fragment within the A2UCOE region on human chromosome 7. ATAC-seq data from PGP1 iPSCs show transcriptionally active regions overlapping with a CpG island between the *HNRNAPA2B1* and *CBX3* genes, suggesting open chromatin in this area. The 0.9 UCOE, isolated from this region, spans 863 bp, including exon 1 of *CBX3* and part of the *HNRNAPA2B1* promoter. B. Schematic of the all-in-one Tet-On plasmid design with and without the 0.9 UCOE. The 0.9 UCOE was placed upstream of the EF1α promoter to drive stable expression of rtTA and selection markers. C. Experimental workflow for generating and analyzing bulk iPSC populations after gene editing. PGP1 iPSCs were edited using CRISPR/Cas9, followed by 5 days of blasticidin selection. Flow cytometry analysis was performed periodically over the 30-day culture period to monitor the percentage of RFP-positive (mScarlet+) cells. D. Fluorescence microscopy images of the 0.9 UCOE and w/o UCOE bulk populations on Days 0 and 30. E. Quantification of RFP-positive cells by flow cytometry in the 0.9 UCOE and w/o UCOE bulk populations over time (n=3). F. Reverse Transcriptase Quantitative PCR (RT-qPCR) analysis of transcription levels for rtTA and mScarlet in bulk populations on Day 0 (n=3). G. RT-qPCR analysis of rtTA and mScarlet transcription levels in bulk populations on Day 30 after 30 days of culture without blasticidin (n=4). H. RT-qPCR analysis of induced *FOXN1* transcription in response to doxycycline on Day 30 after 30 days of culture without blasticidin (n=4). P-values were calculated using a two-tailed unpaired t-test and one-way analysis of variance with Tukey’s honestly significant difference test. Data are presented as mean ± SEM. For detailed data, statistical analyses, and exact p-values, see source data file.

We then evaluated the stability of gene expression in these bulk cell populations post-gene editing (**Figure 2C**), which we note will contain both targeted and untargeted integrants. After a 2-day rest period following gene editing, cell integrants were selected for with 3 µg/mL blasticidin for 5 days, generating gene integrated iPSC colonies. Successful integration of the gene circuits was confirmed by PCR amplification of the newly generated 5’ and 3’ integration junctions at the Rogi1 target region (**Supplemental Figure 1**). Total iPSC colonies were pooled to generate a bulk population. Starting from the day blasticidin selection was completed (Day 0), the bulk cells were cultured for 30 days in medium without blasticidin to permit silencing, and the stability of gene expression downstream of UCOE was evaluated by quantifying the number of RFP-positive (mScarlet+) cells via flow cytometry (**Figure 2C**). Fluorescence microscopy images of the UCOE and w/o UCOE bulk populations on Days 0 and 30 are shown in **Figure 2D**. On Day 0, most colonies and cells were RFP-positive; however, by Day 30 some colonies visibly showed a reduction in RFP presumably due to silencing in both bulk populations. Flow cytometry analysis revealed that on Day 0, the percentage of RFP-positive cells was 46.4% in the 0.9 UCOE bulk population and 41.5% in the w/o UCOE bulk population, showing no significant difference (**Figure 2E**). We attribute the low fraction of RFP-positive cells in the initial population, despite blasticidin selection, to the necessary clump passaging of human iPSCs, which may protect some cells from antibiotic selection. However, by Day 30, the percentage of RFP-positive cells was significantly higher in the 0.9 UCOE population (32.0%) compared to the w/o UCOE population (14.0%) (**Figure 2E**).

Additionally, we evaluated the transcription levels of the introduced genes, rtTA and mScarlet. On Day 0, both rtTA and mScarlet gene transcription levels were approximately 800-fold and 2000-fold higher, respectively, in both the 0.9 UCOE bulk and w/o UCOE bulk groups compared to unmodified parental PGP1 cells. However, there was no significant difference in transcription levels between the 0.9 UCOE and w/o UCOE bulk groups on Day 0 (**Figure 2F**). In contrast, on Day 30, while the expression levels of rtTA and mScarlet decreased compared to Day 0 in the 0.9 UCOE group, they remained approximately 400-fold and 1500-fold higher, respectively. In the w/o UCOE group, a substantial decline in expression levels was observed (**Figure 2G**).

More importantly, the bulk UCOE cells after 30 days still responded to doxycycline administration showing significantly enhanced expression of the target gene, *FOXN1*, whereas by comparison the w/o UCOE group on Day 30 had no response and expression (**Figure 2H**). These results suggest that the A2UCOE promotes long-term stable gene expression in gene-edited iPS cells and helps ensure that cells continue to respond to doxycycline for inducing target genes in the Tet-On system in the absence of selection.

### Establishment of pre-induced thymic epithelial cell line and differentiation into induced thymic epithelial cells (iTEC)

Next, we sought to isolate single-cell-derived clones with the *FOXN1* gene circuit containing the UCOE, for stably making induced thymic epithelial cells (iTEC) (**Figure 3A**). Using CRISPR/Cas9 gene editing, we integrated the 0.9 UCOE-*FOXN1* gene circuit into PGP1 cells and performed blasticidin selection. A total of 24 colonies were picked, and PCR confirmed the correct insertion of the gene circuit in 9 of these cell lines indicating a 37% integration efficiency (**Supplemental Figure 2**). From these, we selected one cell line and established it as our pre-induced thymic epithelial cell line 1 (pre-iTEC1). This cell line exhibited a dense and flat colony morphology indicative of the primed pluripotent state similar to the parental PGP1 cells, and strongly expressed mScarlet fluorescent protein (**Figure 3B**). Reverse Transcriptase quantitative PCR (RT-qPCR) revealed significant expression of rtTA, BSR, and mScarlet compared to PGP1 cells (**Figure 3C**). Additionally, the expression of the pluripotency markers *OCT4*, *NANOG*, and *DNMT3B* were comparable to that in PGP1 cells, indicating that the undifferentiated state was maintained (**Figure 3D**).

**Figure 3.**
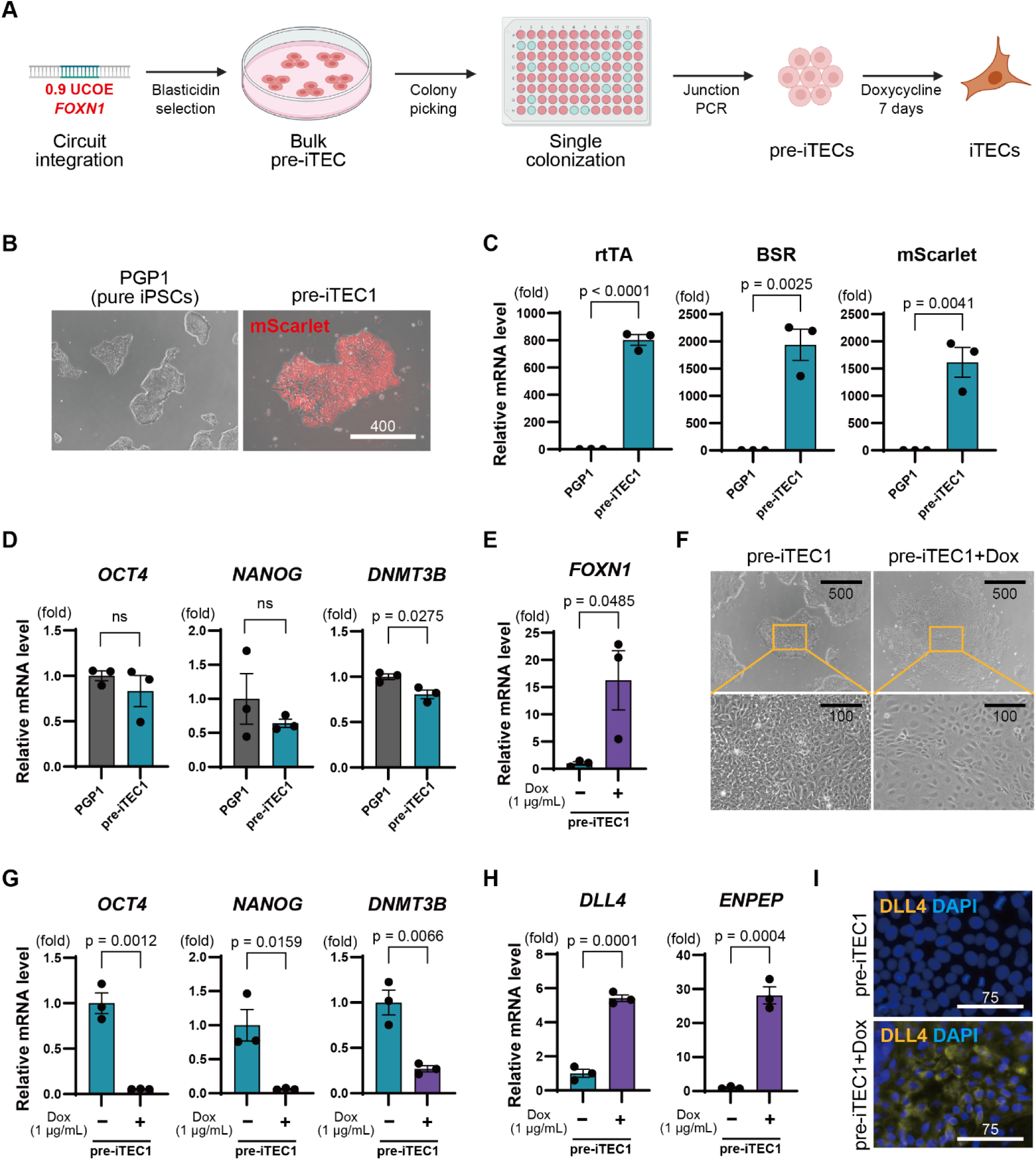
Establishment of pre-induced thymic epithelial cell (pre-iTEC) line and induction of differentiation into thymic epithelial cells (iTEC). A. Experimental scheme: Genetic circuits were introduced into iPSCs to create pre-induced thymic epithelial cells (pre-iTEC). Induction into induced thymic epithelial cells (iTEC) was achieved by administering 1 µg/mL doxycycline for 7 days. B. Representative images of the created pre-iTEC1 and PGP1 cells. The RFP signal indicates the expression of the mScarlet gene. C. RT-qPCR comparison of the introduced genes rtTA, BSR, and mScarlet between groups (n=3). D. RT-qPCR comparison of undifferentiated marker genes *OCT4*, *NANOG*, and *DNMT3B* in PGP1 cells and pre-iTEC1 (n=3). E. RT-qPCR comparison of *FOXN1* gene expression in pre-iTEC1 and pre-iTEC1 treated with doxycycline groups (n=3). F. Morphological changes in pre-iTEC1 upon doxycycline administration. G. RT-qPCR comparison of undifferentiated markers in pre-iTEC1 with and without doxycycline treatment (n=3). H. Comparison of TEC markers *DLL4* and *ENPEP* (Ly51) in pre-iTEC1 upon doxycycline treatment (n=3). I. Representative image of immunofluorescence staining for DLL4 in pre-iTEC1 treated with doxycycline. P values were calculated using a two-tailed unpaired t-test. The data are presented as mean ± SEM. For detailed data, statistical analyses, and exact p-values, see source data file.

Next, we evaluated the forward programming potential of pre-iTEC1 iPSCs into iTECs. Upon treatment with doxycycline (1 µg/mL, 24 hours), *FOXN1* expression was significantly induced in the doxycycline-treated group, as confirmed by RT-qPCR (**Figure 3E**). Moreover, when 1 µg/mL of doxycycline was administered for 7 days in B8-iPSC media, which enforces the pluripotent state, the morphology of the cells changed from the dense colony-like appearance characteristic of iPSCs to a looser and elongated form (**Figure 3F**, **Supplemental Figure 3**), indicating that pre-iTEC cells underwent morphological changes upon doxycycline treatment even in the presence of media meant to maintain pluripotency. The expression of the pluripotency markers *OCT4*, *NANOG*, and *DNMT3B* was significantly decreased in the doxycycline-treated group (**Figure 3G**). Furthermore, we observed an increase in expression of the key TEC markers *DLL4* and *ENPEP* (Ly51) in the doxycycline-treated group (**Figure 3H**). Immunostaining further revealed strong DLL4 protein expression in the doxycycline-treated group (**Figure 3I**). These results demonstrate that pre-iTEC1 iPSCs reprogram into iTECs upon *FOXN1* induction alone.

### 0.9 UCOE induces gene leakage and unintended differentiation of iPS cells

However, we observed *FOXN1* gene leakage in the pre-iTEC1 cell line even in the absence of doxycycline (**Figure 4A**). When pre-iTEC1 was cultured without doxycycline for 30 days, we noted a significant decrease in the pluripotency markers *OCT4*, *NANOG*, and *DNMT3B*, indicating unintended differentiation, whereas PGP1 cells maintained their pluripotency over the same 30-day culture period (**Figure 4B**). Furthermore, after long-term culture for 90 days without doxycycline, we observed increased transcription of the TEC-specific markers *DLL4* and *ENPEP* (Ly51), despite the absence of doxycycline (**Figure 4C**).

**Figure 4.**
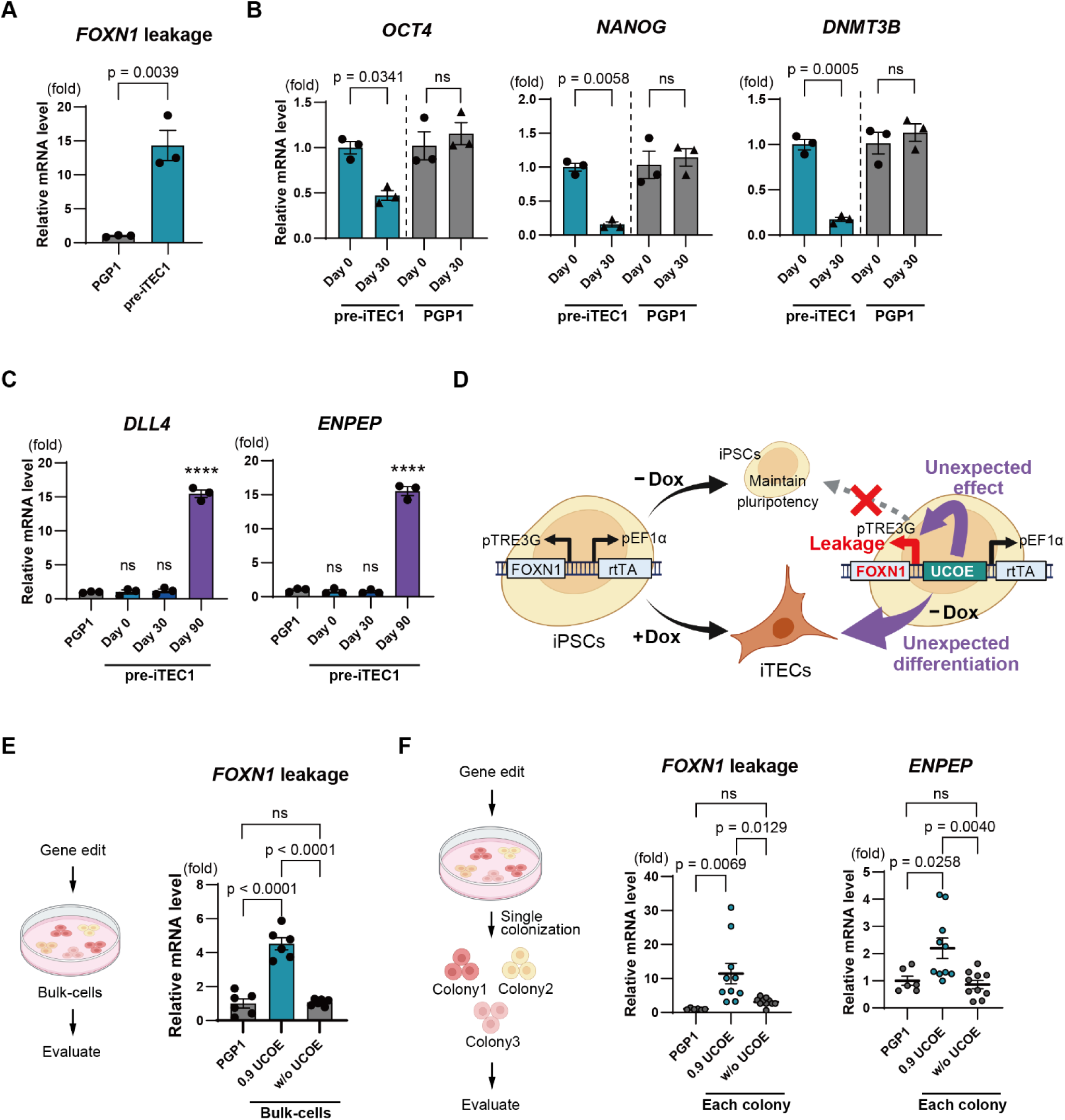
0.9 UCOE induces gene leakage and unintended differentiation of iPS cells. A. Analysis of *FOXN1* gene leakage in pre-iTEC1 and PGP1 without doxycycline treatment (n=3). B. RT-qPCR comparison of undifferentiated marker genes *OCT4*, *NANOG*, and *DNMT3* in pre-iTEC1 and PGP1 cultured without doxycycline for 30 days (n=3). C. RT-qPCR comparison of TEC-specific markers *DLL4* and *ENPEP* (Ly51) in pre-iTEC1 during 90 days culture without doxycycline (n=3). D. Hypothetical model. iPSCs with drug-inducible circuits are expected to maintain pluripotency in the absence of doxycycline. However, 0.9 UCOE may cause *FOXN1* leakage, leading to unintended differentiation of iPSCs into iTECs. E. Analysis of *FOXN1* leakage in bulk cell populations of the 0.9 UCOE and w/o UCOE group by RT-qPCR (n=3). F. *FOXN1* leakage and *ENPEP* (Ly51) expression analysis in single-cell-derived colonies of 0.9 UCOE group and w/o UCOE group by RT-qPCR (n=10). P values were calculated using a two-tailed unpaired t-test and one-way analysis of variance with Tukey’s honestly significant difference test. Data are presented as mean ± SEM. For detailed data, statistical analyses, and exact p-values, see the source data file.

These results led us to formulate a hypothesis regarding the role of the 0.9 UCOE (**Figure 4D**). Generally, iPSCs with drug-inducible circuits are expected to maintain pluripotency in the absence of the small molecules like doxycycline. However, we hypothesize that the 0.9 UCOE may have contributed to unexpected *FOXN1* expression leakage, leading to unintended differentiation of pre-iTEC iPSCs into iTECs. To test this hypothesis, we integrated both 0.9 UCOE and w/o UCOE gene circuits into PGP1 cells, generated new bulk groups after blasticidin selection, and evaluated the level of *FOXN1* leakage in each group. The w/o UCOE group showed no significant difference in *FOXN1* leakage compared to the control PGP1 cells, while the UCOE group exhibited significantly higher *FOXN1* leakage in the absence of doxycycline (**Figure 4E**). Next, we compared single-cell-derived clones from both groups. Ten colonies each were picked from both the 0.9 UCOE and w/o UCOE groups, and precise integration at Rogi1 was confirmed by PCR (**Supplemental Figure 4**). When comparing *FOXN1* leakage among these clones, no significant *FOXN1* leakage was observed in the w/o UCOE colonies. In contrast, the 0.9 UCOE colonies displayed significant *FOXN1* leakage and greater heterogeneity in absolute expression from clone to clone (**Figure 4F**). Moreover, the 0.9 UCOE colonies showed elevated expression of the TEC marker *ENPEP*, even shortly after cell line establishment (**Figure 4F**). These findings indicate that 0.9 UCOE contributes to gene leakage in the Tet-On system, leading to unexpected differentiation of iPS cells.

Additionally, we cultured the 0.9 UCOE bulk group and the w/o UCOE bulk group for 30 days in the presence of blasticidin but without doxycycline to evaluate morphological changes (**Supplemental Figure 5**). As described in **Figure 3F**, the w/o UCOE bulk group maintained a dense colony-like appearance, characteristic of iPS cells. In contrast, some cells in the 0.9 UCOE bulk group exhibited a looser and elongated form, indicating that *FOXN1* leakage driven by the 0.9 UCOE resulted in unexpected differentiation, as evidenced by morphological changes.

### Evaluation of gene stability and gene leakage using various lengths of A2UCOE fragments

Next, we generated several A2UCOE fragments to investigate whether longer or shorter forms could reduce gene leakage while enhancing expression stability. Previous studies have examined various lengths of A2UCOE fragments and their abilities to promote long-term stable expression; however, no studies to date have evaluated A2UCOE in terms of gene leakage. Using the UCSC Genome Browser, we identified the precise location of A2UCOE in the human genome along with information on nearby promoters and enhancers (**Supplemental Figure 6**). Based on this information, we designed three new UCOE fragments (**Figure 5A**, **Supplemental Table 1**). The 1.3 UCOE fragment, 1,337 bp in length, contains the untranslated region (UTR) and exon 1 of *HNRNAPA2B1*, along with EH38E2541925, a promoter-like element associated with *HNRNAPA2B1*, extending to include EH38E2541926, a promoter region and exon 1 of *CBX3*. The 0.7 UCOE fragment, 749 bp in length, does not include the *HNRNAPA2B1* exon 1 or promoter region but contains the *CBX3* exon 1 and promoter region, as well as the intergenic region between EH38E2541925 and EH38E2541926. The CBX UCOE fragment is shorter, at 547 bp, and includes only the *CBX3* promoter region and exon 1. These newly designed fragments, along with the previously used 0.9 UCOE and w/o UCOE fragments, were each incorporated into the *FOXN1* drug-inducible gene circuit. These gene circuits were integrated into the Rogi1 safe harbor site in PGP1 cells using CRISPR/Cas9 gene editing, followed by a 5-day blasticidin selection to create bulk populations for each UCOE variant. The day on which the blasticidin selection was completed was defined as Day 0, and the cells were cultured in the absence of blasticidin for 30 days to evaluate gene expression.

**Figure 5.**
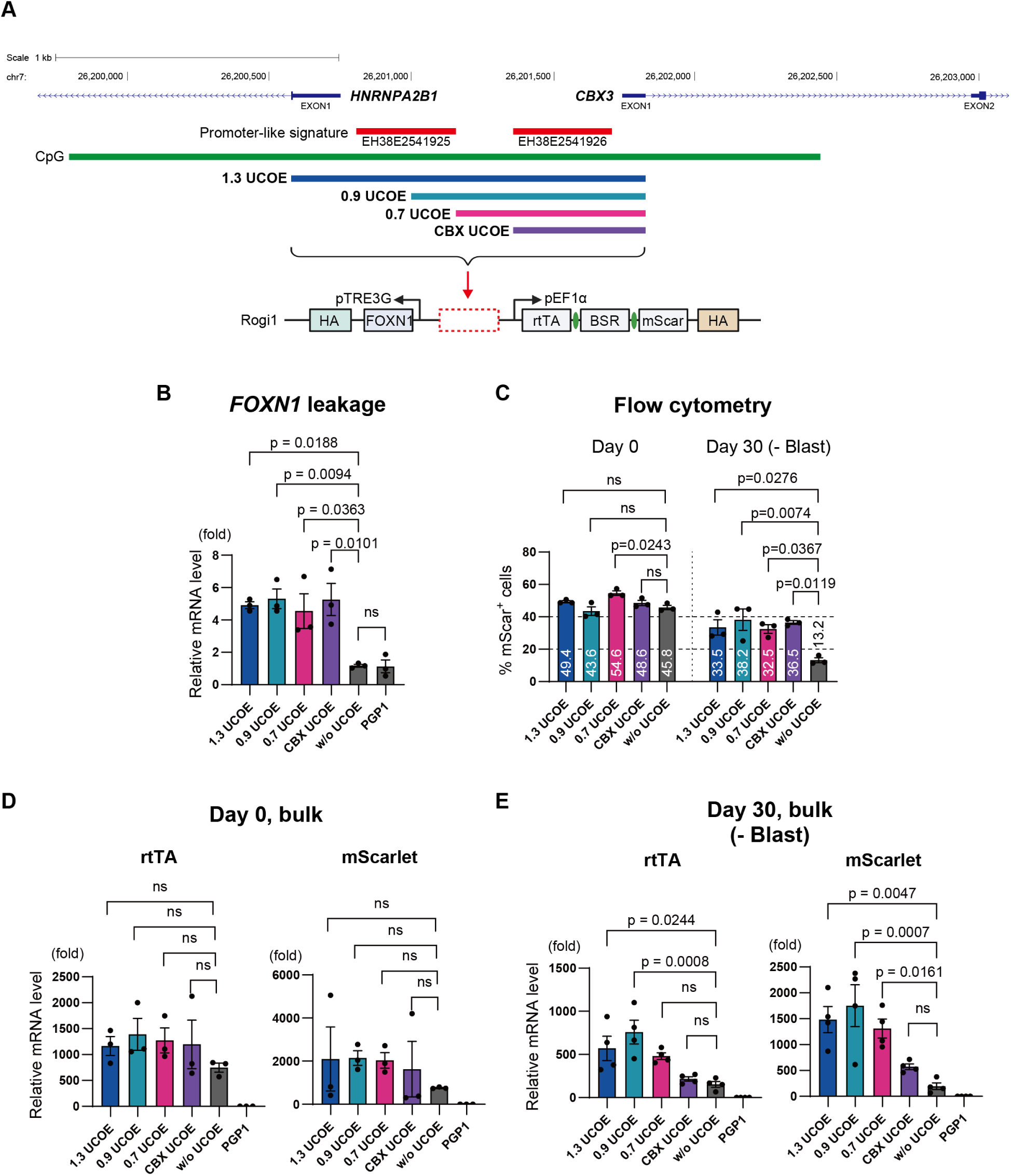
Evaluation of gene stability and gene leakage using various lengths of A2UCOE fragments. A. Design of different A2UCOE fragments. The 1.3 UCOE (1,337 bp) includes the untranslated region (UTR) and exon 1 of *HNRNAPA2B1*, as well as the promoter-like regions EH38E2541925 and EH38E2541926 extending to include the *CBX3* exon 1. The 0.9 UCOE (863 bp) included the part of the *HNRNAPA2B1* promoter and extending to exon 1 of *CBX3*. The 0.7 UCOE (749 bp) contains only the *CBX3* exon 1, promoter region, and intergenic sequence. The CBX UCOE (547 bp) includes only the *CBX3* promoter region and exon 1. These fragments were incorporated into the *FOXN1* Tet-On inducible gene circuit and integrated into the Rogi1 site of PGP1 iPSCs. B. *FOXN1* gene leakage in each UCOE group assessed on Day 0 by RT-qPCR (n=3). C. Flow cytometry analysis of RFP-positive cells in each UCOE group on Day 0 and Day 30 (n=3). D. Transcription levels of rtTA and mScarlet on Day 0 in each UCOE group by RT-qPCR (n=3). E. Transcription levels of rtTA and mScarlet on Day 30 in each UCOE group after 30 days of culture without blasticidin by RT-qPCR (n=3). P values were calculated using one-way analysis of variance with Tukey’s honestly significant difference test. The data are presented as mean ± SEM. For detailed data, statistical analyses, and exact p-values, see source data file.

We then compared *FOXN1* gene leakage and the stability of downstream gene expression across different UCOEs. First, we assessed *FOXN1* leaky expression on Day 0 and found that all UCOE fragments exhibited significant gene leakage at comparable levels relative to the w/o UCOE group (**Figure 5B**). Next, we quantified RFP-positive (mScarlet+) cells by flow cytometry. On Day 0, the percentages of RFP-positive cells were 49.4% for 1.3 UCOE, 43.6% for 0.9 UCOE, 54.6% for 0.7 UCOE, and 48.6% for CBX UCOE. Among these, only the 0.7 UCOE group showed a significant difference compared to the w/o UCOE control group, while the other UCOE fragments did not show any significant differences. By Day 30, the percentages were 33.5%, 38.2%, 32.5%, and 36.5% for 1.3 UCOE, 0.9 UCOE, 0.7 UCOE, and CBX UCOE, respectively. All UCOE fragments showed a significant difference compared to the w/o UCOE control group; however, there were no significant differences among the four UCOE fragments (**Figure 5C**, **Supplemental Figure 7**).

We also evaluated the transcription levels of the selection markers, rtTA and mScarlet. On Day 0, transcription levels of rtTA and mScarlet were comparable across all UCOE fragments, with no significant differences relative to the w/o UCOE control group (**Figure 5D**). However, by Day 30, transcription levels of rtTA were significantly maintained in the 1.3 UCOE and 0.9 UCOE groups compared to the w/o UCOE group. For mScarlet, transcription levels were significantly maintained in the 1.3 UCOE, 0.9 UCOE, and 0.7 UCOE groups compared to the w/o UCOE group. In the CBX UCOE group, transcription levels of both rtTA and mScarlet had decreased to levels comparable to those in the w/o UCOE group (**Figure 5E**).

These results indicate that an A2UCOE length of at least 0.7 kb positively influences the maintenance of transcription levels, whereas the CBX-only UCOE fragment is insufficient to sustain transcription. However, when assessing the RFP-positive cell number via flow cytometry, no significant differences were observed between UCOE fragments of 0.7 kb or longer and the shorter CBX UCOE. Nevertheless, examining fluorescence output revealed that the intensity peak for the CBX-UCOE shifted significantly to the left compared to the peaks of the other three UCOE fragments, suggesting that the CBX UCOE results in a higher number of weakly positive RFP cells (**Supplemental Figure 8**). Additionally, when doxycycline was administered to the CBX UCOE group after 30 days of culture, *FOXN1* transcription levels were significantly higher compared to the w/o UCOE group and comparable to the 0.9 UCOE group (**Supplemental Figure 9**). This finding suggests that, despite the lower transcription and protein expression levels of rtTA in the CBX UCOE group, the inducible gene expression function was stably maintained over an extended period. Overall, these results demonstrate that any UCOE fragment can sustain inducible gene expression function over time compared to the condition without UCOE.

### SV40 poly-A spacer sequences mitigate gene leakage caused by A2UCOE

Finally, we sought to address the gene leakage caused by A2UCOE. Since UCOE sequences facilitate open chromatin in bidirectional gene clusters, we considered the possibility that it stabilizes RNA polymerase II elongation in both directions, despite the inducible TRE promoter lacking bound rtTA activating proteins. Whereas the UCOE contains a high density of CpG dinucleotides thought to prevent silencing, AT-rich sequences are thought to condense chromatin and resist protein binding, and are often found in termination sequences, matrix attachment regions, and replication origins. We hypothesized that inserting an AT-rich spacer sequence between the A2UCOE and TRE promoter might reduce the leaky gene expression. To test this hypothesis, we introduced several AT-rich spacer sequences between A2UCOE and TRE-*FOXN1*. We designed random 238 bp spacer sequences with evenly distributed AT content of 35%, 50%, and 65%, labeled AT35, AT50, and AT65, respectively (**Figure 6A**, **Supplemental Table 2**). Additionally, we included a 238 bp spacer sequence with 65% AT content that contained the 122 bp termination sequence SV40 poly-A (SV40), which includes homopolymeric A and T tracts known to terminate transcription by forming hairpin structures and recruiting termination proteins.

**Figure 6.**
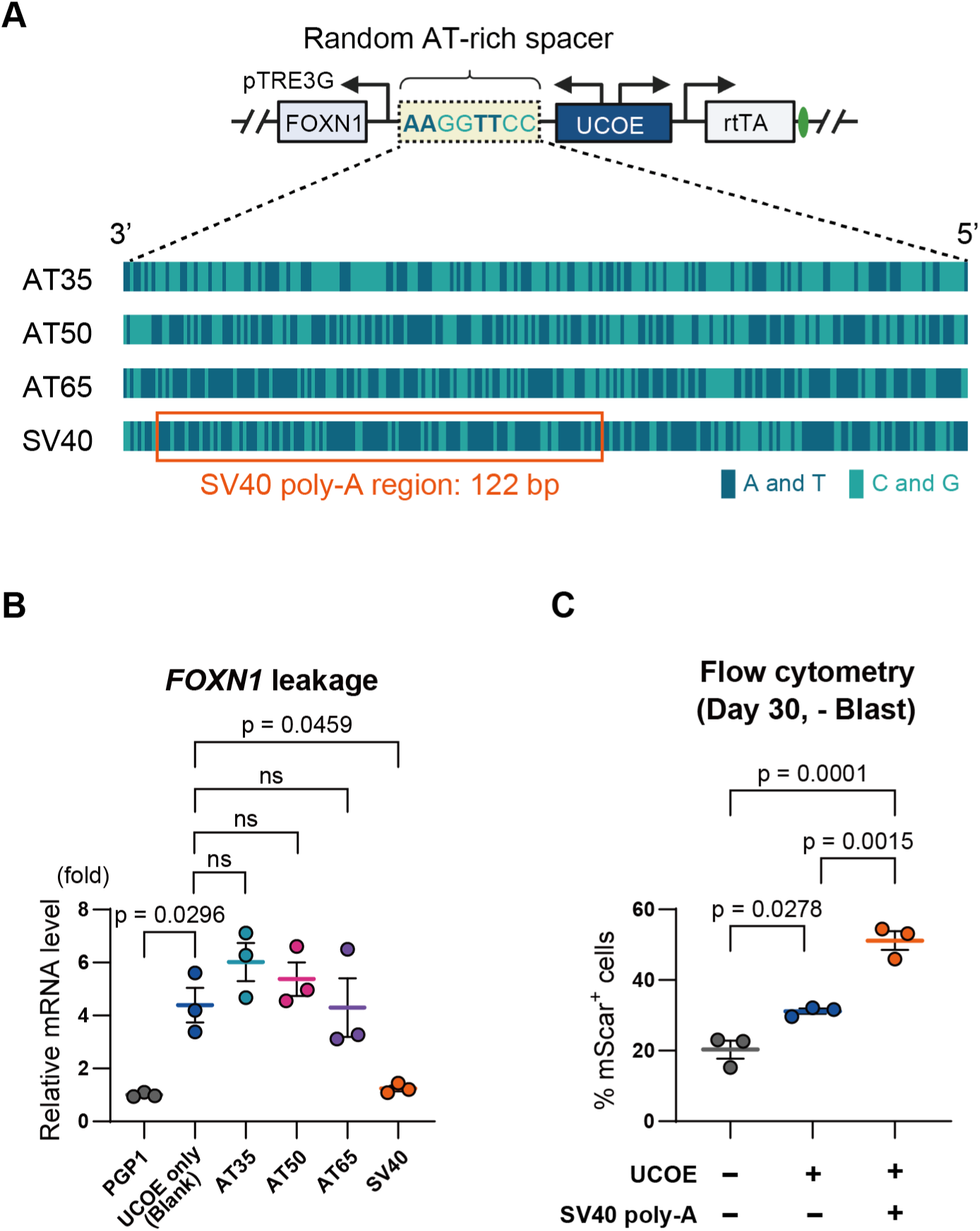
SV40 poly-A spacer sequences mitigate gene leakage caused by A2UCOE. A. Schematic of gene circuit architecture with various AT-rich spacer sequences between A2UCOE and the TRE-*FOXN1* promoter. Spacer sequences were designed randomly with 238 bp lengths and varied AT content: about 35% (AT35), 50% (AT50), and 65% (AT65), as well as a 65% AT-rich spacer containing the 122 bp SV40 poly-A termination sequence (SV40). The dark green and light green bars represent the sequence composition, with A and T shown in dark green and C and G in light green. The SV40 poly-A sequence is highlighted with an orange box. B. Quantification of *FOXN1* gene leakage in bulk cell populations following the integration of gene circuits with different AT-rich spacers (n=3). C. Flow cytometry analysis of RFP-positive (mScarlet+) cells in the SV40 poly-A spacer group after 30 days of culture without blasticidin (n=3). P values were calculated using one-way analysis of variance with Tukey’s honestly significant difference test. The data are presented as mean ± SEM. For detailed data, statistical analyses, and exact p-values, see source data file.

These gene circuits were integrated into the Rogi1 site in PGP1 cells as before, followed by a 5-day blasticidin selection to generate bulk populations. We then evaluated *FOXN1* gene leakage in these populations. As shown in **Figure 6B**, the SV40 spacer significantly reduced *FOXN1* leakage to nearly the same level as the control group. In contrast, the AT35, AT50, and AT65 spacers did not show a significant reduction in leakage compared to the UCOE alone, suggesting that a 238 bp separation alone is insufficient to mitigate leakiness, nor does AT content alone seem to influence gene leakage.

Next, we tested whether this new gene circuit architecture containing the SV40 poly-A spacer preserved the original UCOE benefits of anti-silencing and enhanced transcriptional stability. As before, cells were cultured without blasticidin for 30 days, and RFP-positive (mScarlet+) cells were quantified by flow cytometry. Even in the group containing the SV40 poly-A spacer, the number of RFP-expressing cells was maintained on Day 30 compared to the w/o UCOE control group (**Figure 6C**). Moreover, the SV40 poly-A group also showed significantly better maintenance of RFP-positive cell numbers compared to the UCOE-only group (**Figure 6C**). These results demonstrate that the SV40 poly-A termination sequence effectively reduces gene leakage caused by A2UCOE without interfering with its original anti-silencing and transcription-stabilizing benefits.

## Discussion

In this study, we demonstrated that the A2UCOE causes unintended gene leakage when paired with an all-in-one Tet-On system driving the master transcription factor *FOXN1*, leading to premature differentiation of iPSCs. The leaky expression is best explained by a model in which the open chromatin promoted by A2UCOE enables binding and elongation of RNA polymerase II (RNAPII) in both transcriptional directions without the aid of activating factors (**Figure 7**). We show that the leaky gene expression can be mitigated by inserting a spacer containing poly-A termination sequence, but not necessarily with AT-rich tracts alone. While *in vitro* studies show that AT-rich tracts cause RNAPII pausing and termination, sometimes independent of specific protein factors (40–42), the specific poly-A termination signal was necessary to eliminate gene leakage. This likely means that recruitment of termination factors like poly-A polymerase and poly-A binding protein (PABPC) at canonical poly-A signal sequences are necessary for 3’-end processing and termination of basal transcription (43). In fact, the SV40 spacer appeared to enhance the anti-silencing effect of the UCOE, which may perhaps mean that epigenetic silencing senses aspects of where RNAPII is located. We recommend then that when assembling synthetic gene circuits with both inducible and constitutive promoters with the A2UCOE, that an SV40 poly-A sequence should be introduced between transcriptional units to block premature RNAPII elongation. Further work might focus on the minimal number of spacer sequence elements that would confer these effects similar to the SV40 terminator.

**Figure 7.**
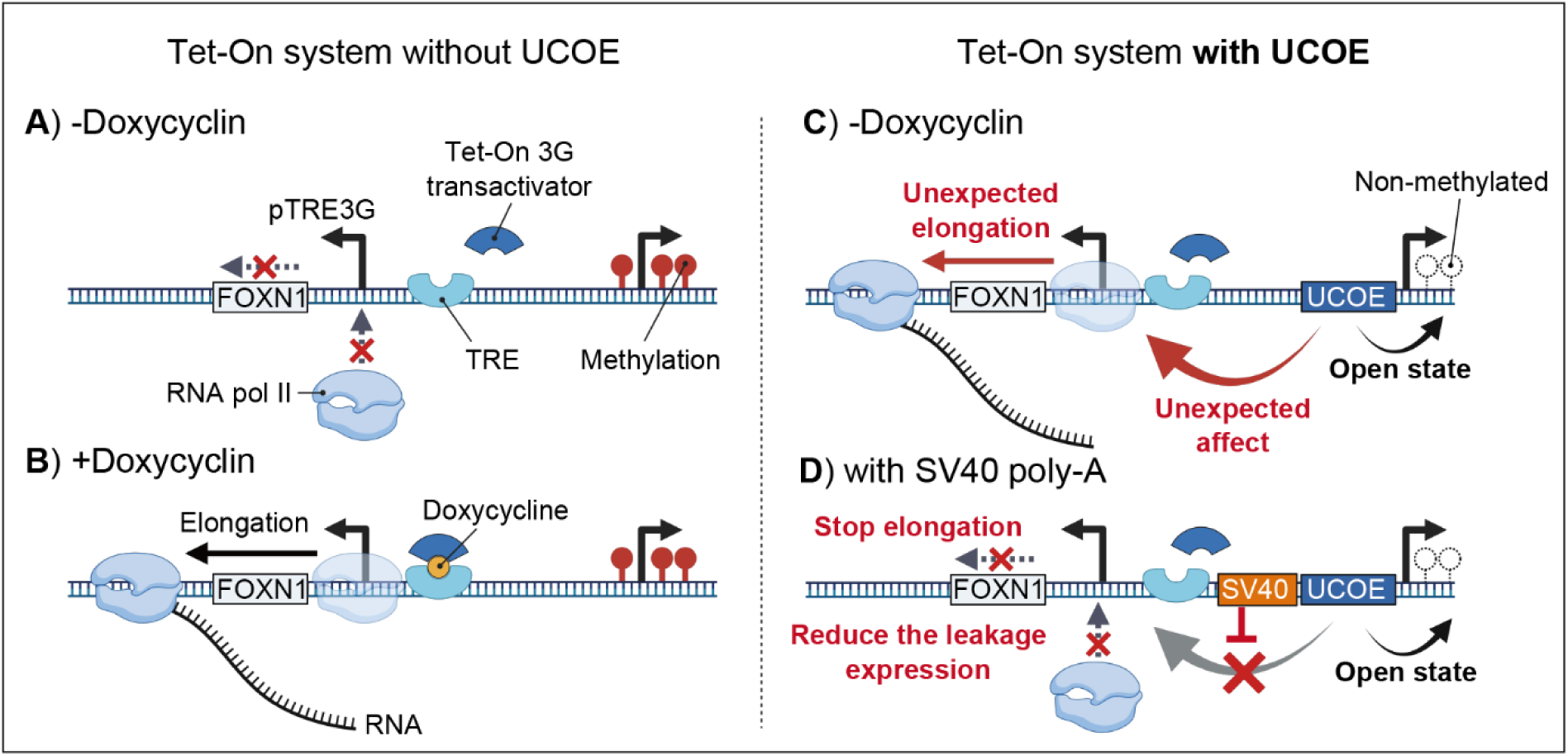
Mechanism of gene leakage in the Tet-On 3G system with A2UCOE. A. In the absence of doxycycline and A2UCOE, the Tet-On 3G transactivator does not bind to the tetracycline response element (TRE), and RNA polymerase II (RNA pol II) elongation does not initiate. The integrated gene will be silenced through methylation. B. In the presence of doxycycline, the Tet-On 3G transactivator binds to the TRE, initiating RNA pol II elongation and resulting in transcription. C. In an environment with A2UCOE but without doxycycline, the A2UCOE maintains the sense-strand constitutively expressed genes (rtTA, BSR, mScarlet) in an open chromatin state, preventing methylation and silencing. However, A2UCOE also exerts an unexpected effect on RNA pol II on the anti-sense strand, leading to unexpected elongation through the TRE promoter causing gene transcription even in the absence of doxycycline. D. When the SV40 poly-A sequence is present, it blocks the influence of A2UCOE on RNA pol II, stopping elongation and thereby reducing gene leakage.

We also introduce a user-friendly all-in-one Tet-On system that adopts a compact (5 kb) bidirectional design, where the EF1α promoter and TRE3G promoter are oriented in opposite directions. The A2UCOE with the SV40 poly-A terminator has minimal basal transcription, and resists silencing better than without. Compact Tet-On systems are particularly useful for lentiviral vector genomic integrations, which have a payload size limitation of 8–10 kb (11,12). However, our system utilizes site-specific integration via CRISPR/Cas9 assisted homology-directed repair into a recently characterized safe-harbor site Rogi1 (38). While this safe-harbor site was reported to be better than other sites, it is still subject to epigenetic silencing, at least in human iPSCs. Nevertheless, our all-in-one system is easily deployed to other genomic sites. Introducing homology arms to new genomic sites requires only PCR of 5’ and 3’ ∼450 bp homology arms appended with BsaI sites. This facilitates facile Golden Gate assembly in a one-pot reaction. Genomic integration is then assisted with a CRISPR/Cas9 gRNA directed at the site, which can either be expressed from a plasmid or delivered as an *in vitro* assembled ribonucleoprotein. We typically obtain numerous iPSC clones with a 6 kb construct, and anecdotally, similar numbers of clones for constructs up to at least 10 kb.

The system is particularly useful when sustained inducible expression is required, but without any leaky basal expression. This was particularly important when we expressed the pioneer transcription factor *FOXN1*, which was powerful enough to induce a thymic epithelial cell-like identity from leaky expression alone and in pluripotency media. Other direct reprogramming pioneer factors have the same ability to promote new cell identity alone. Transcription factors such as *MYOD1* (muscle) (44), *NGN2* (neurons) (45), and *ETV2* (endothelial) (46), to name a few, also may cause premature differentiation in iPSCs if expression is not tightly controlled, at least when containing an A2UCOE. As mammalian synthetic biology progresses towards more applied applications where complex gene circuits may be required, it will be important to further understand the rules by which transcription occurs based on the arrangement and architecture of DNA regulatory elements. The importance of spacing and orientation has been recently highlighted in modeling work of circuits similarly-sized to our system showing that transcriptional orientation can induce negative effects due to DNA supercoiling (47).

One limitation of our study is that the impact of A2UCOE on basal expression was only evaluated in the Tet-On system, and we did not test other inducible circuits. Recently, several new mammalian small molecule-inducible systems, such as the Cumate and synthetic zinc finger transcription regulators (synZiFTR) systems have been developed for controlled gene expression (48,49). It is possible that UCOE-induced gene leakage may behave differently in these systems due to differences in the promoter architecture. However, the Tet-On system is widely used and has undergone numerous improvements, making it one of the least leaky systems available. Despite using an improved Tet-On 3G system known for minimal leakage because it lacks transcription factor binding sites, we still observed UCOE-induced gene leakage. This suggests that UCOE could potentially have a negative impact on gene leakage in other drug-inducible circuits as well.

In summary, this study revealed that the A2UCOE can cause unintended gene leakage in bidirectional inducible gene circuits, leading to premature differentiation of iPS cells. This finding highlights a potential pitfall in the design of gene circuits for direct reprogramming, particularly when maintaining pluripotency is critical, such as in iPS and ES cells. This is also of extreme importance for therapeutic payloads where sustained expression and tight regulation are necessary for safety. Future designs of gene circuits should consider the requirements needed for DNA found in between transcription elements, as is already naturally found in mammalian genomes.

## Acknowledgements

This work was supported by National Institute of Allergy and Infectious Diseases grant DP2AI154417 to DMT.

## Author Contributions

Performed Research: TY, SFP, JA, DEA. Experimental Design: TY, DMT. Analyzed Data: TY, SFP. Wrote Manuscript: TY, DMT. Funding: DMT.

## Materials and Method

### ATAC-seq analysis using external database data

ATAC-seq data was obtained from series GSE170231, which hosts publicly available chromatin accessibility profiles (35). Fastq data sets for ATAC-seq on human cell line PGP1 were downloaded and integrated into our analysis. Following data retrieval, quality control checks were conducted, including assessments of read depth and signal-to-noise ratio to ensure data integrity. Sequencing reads were aligned to the human reference genome GRCh38 using Bowtie 2. Peak calling was performed using IGV.

### Cell culture

Human iPSC line PGP1 was obtained from George Church’s laboratory. Cells were grown at 37°C at 5% CO2 and ambient (∼19%) oxygen in B8 iPSC medium (50), which contains DMEM/F12 (Corning, NY, USA), 200 µg/mL of L-Ascorbic acid 2-phosphate, 5 µg/mL of human insulin (Gibco), 5 µg/mL of human transferrin (InVitria), 20 ng/mL of sodium selenite, 40 ng/mL of fibroblast growth factor 2-G3 (FGF2-G3) (made at Northwestern University core facility), 0.1 ng/mL of neuregulin 1 (NRG1) (Peprotech), and 0.1 ng/mL of transforming growth factor beta-3 (TGFb3) (Peprotech), on plates coated with Cultrex (R&D systems, Inc, MN, USA), and they were grown until achieving about 70% confluence. Passaging of cells was performed by clump passaging using 0.5 mM EDTA, and with Accutase (Innovative Cell Technologies) when single cell dissociation was required. Cells were assayed regularly for mycoplasma.

### Construction of Genetic Circuits

The bidirectional Tet-On plasmid system was assembled from DNA fragments obtained from different source plasmids available from Addgene or by chemical synthesis from Integrated DNA Technologies (IDT, USA). Cloning was performed using a mixture of standard restriction enzyme cloning and Gibson assembly. The plasmid backbone is based on the yeast vector pRS416 (Boeke laboratory) and contains E. coli replication elements including the ampicillin resistance marker. Various lengths of UCOE fragments, including a 0.9 kb-UCOE, were either synthesized by IDT or PCR amplified from human genomic DNA. The *FOXN1* gene was extracted from the plasmid provided by Addgene (Addgene#101445). These genes were assembled into genetic circuits using Gibson assembly with E. coli strains (TOP10, EPI300, and dam-/dcm-competent cells) as well as yeast-based assembly. The constructed gene circuits were subsequently verified for sequence accuracy through full-length sequencing performed by Plasmid-EZ service (Azenta, USA).

### Gene Integration

For targeted gene integration into the Rogi1 site on chromosome 1, we designed guide RNAs (gRNAs) to specifically cleave this region. These gRNAs were expressed using a plasmid based on the PX459 vector (Addgene#62988). Next, homology arms (HAs) of approximately 450 bp each were designed for the 5’ and 3’ sides of the Rogi1 cleavage site and amplified by PCR to create DNA fragments. For one-pot Golden Gate cloning, we combined 10 ng of the 5’ HA, 10 ng of the 3’ HA, 200 ng of the *FOXN1*-containing all-in-one Tet-On plasmid, 1 μL of BsaI, 1 μL of T4 DNA ligase, 1 μL of T4 ligation buffer (10x), and 1 μL of ATP, adjusting the total volume to 10 μL with distilled water. The reaction mixture was incubated overnight at 37°C. Then, to prepare for homology-directed repair (HDR), the plasmid containing the homology arms was linearized using restriction enzymes (Kpn1 and Nhe1) at the ends of the homology arms. These sequences of the gRNAs and homology arms are provided in **Supplemental Table 2**. The linearized DNA, along with the plasmid expressing the gRNAs, was introduced into cells via nucleofection using the P3 Primary Cell 4D-Nucleofector kit (Lonza). Following nucleofection, cells were subjected to selection with 3 µg/mL of blasticidin to ensure the survival of only those cells that successfully integrated the desired genetic constructs. After the selection process, the accuracy of the genetic editing was verified using junction PCR, a technique designed to confirm the presence of the gene-editing event at the intended genomic location. The primer sequences used for this PCR are listed in **Supplemental Table 3**.

### Cell differentiation protocol

Once cell confluency reached approximately 40-50%, the culture medium was switched to differentiation medium. The differentiation medium consisted of Advanced DMEM supplemented with 60 mg/L of 2-ascorbic acid 2-phosphate, 10 mL/L of GlutaMAX™, and 15 mL/L of 1M HEPES (Gibco). To induce differentiation, 1 µg/mL of doxycycline was added to the medium. Cells were then cultured in this condition for a period of 7 days to promote differentiation.

### Counting RFP-positive cells by flow cytometry

Cells were detached using Accutase and fully dissociated into a single-cell suspension. The cell count was determined using the Countess 3 (ThermoFisher), and the cells were resuspended at a concentration of 5,000 cells/µL in 1 mM EDTA-PBS solution. The suspension was then passed through a 35 µm filter to ensure complete dissociation into individual cells. Flow cytometry was performed using a BD Accuri C6 flow cytometer (Becton Dickinson Company, NJ, USA). A total of 10,000 cells were analyzed per sample, and the number of RFP-positive cells was measured using the PE fluorescence channel. The data were analyzed using Floreada.io.

### RNA extraction and cDNA synthesis

RNA was extracted from cells at 80-90% confluence using the Quick-RNA Miniprep Kit (Zymo Research, CA, USA), following the manufacturer’s protocol. The quantity of extracted RNA was then measured using a NanoDrop spectrophotometer (Thermo Fisher Scientific, MA, USA). Subsequently, reverse transcription via MMLV-RT was performed using ABScript Neo RT Master Mix (ABclonal, Inc., MA, USA) to synthesize cDNA according to the instructions provided by the manufacturer.

### Quantitative real-time PCR (qPCR) analysis

Quantitative PCR was performed using the Forget-Me-Not™ EvaGreen® qPCR Master Mix manufactured by Biotium (CA, USA). All samples were prepared according to the manufacturer’s instructions. To ensure reproducibility and accuracy, each sample was measured in technical duplicates or triplicates, and the results were averaged. The expression levels of the target genes were normalized to the housekeeping gene *RPS29* using the Ct values. Relative gene expression was calculated using the ΔΔCT method. The list of primers used is provided in **Supplemental Table 4**.

### FOXN1 Leakage Analysis

We evaluated *FOXN1* gene leakage using qPCR. The *FOXN1* coding sequence used in this study was derived from a plasmid obtained from Addgene (Addgene #101445). To analyze gene leakage, we designed three primer sets specific to the synthetic *FOXN1* gene using Primer-BLAST (https://www.ncbi.nlm.nih.gov/tools/primer-blast/) (**Supplemental Figure 10**). qPCR was performed on three experimental groups: parental PGP1 cells, bulk iTEC cells containing the integrated 0.9 UCOE-FOXN1 gene circuit, and bulk cells with the integrated 0.9 UCOE-FOXN1 gene circuit treated with doxycycline. Among the three primer sets, Primer Set 1, which produced the least variation in Ct values and the most consistent amplification curve, was selected and used throughout this study.

### Immunofluorescence staining

For immunofluorescence, cultured cells were first aspirated to remove the supernatant, and then washed with phosphate-buffered saline (PBS). Cells were fixed by adding 4% paraformaldehyde (PFA) and incubated for 30 minutes at room temperature. After fixation, the cells underwent three washes with PBS. Blocking was performed using 1% bovine serum albumin (BSA) in PBS with 0.1% Tween 20 (PBST) for 30 minutes to prevent non-specific binding. DLL4 polyclonal antibody (#PA5-97664, Thermo Fisher Scientific, MA, USA) was diluted at 1:200 in 1% BSA in PBST and applied to the cells, followed by overnight incubation at 4°C. After incubation, cells were washed three times with PBS. Goat anti-Rabbit IgG (H+L) Cross-Adsorbed Secondary Antibody, Alexa Fluor™ 488 (Thermo Fisher Scientific, MA, USA) was then diluted at 1:2000 in 1% BSA in PBST and applied to the cells. The cells were incubated for 1 hour in the dark at room temperature, followed by washing with PBS. Stained cells were imaged using the EVOS™ M7000 Imaging System (Thermo Fisher Scientific, MA, USA).

### Design and Creation of Various A2UCOE Fragments

Using the UCSC Genome Browser (https://genome.ucsc.edu/), we obtained information on the A2UCOE region, including neighboring gene sequences, promoters, and enhancers (**Supplemental Figure 6**). Based on this information, we designed the 1.3 UCOE, 0.9 UCOE, 0.7 UCOE, and CBX UCOE fragments. Gene fragments were synthesized by Integrated DNA Technologies (IDT, USA). The sequences of each synthesized A2UCOE fragment are provided in **Supplemental Table 5**.

### Design and Creation of Random AT-Rich Spacer Sequences

To create random sequences, we used ChatGPT-4 to generate 238 bp sequences with approximately 35%, 50%, and 65% AT content. These gene fragments were synthesized by Integrated DNA Technologies (IDT, USA). The sequence information for each spacer is provided in **Supplemental Table 1**.

### Statistical analysis

Statistical analyses were conducted using GraphPad Prism 10.2.2 software. For comparisons between two groups, unpaired t-tests were used to determine statistical significance. For analyses involving three or more groups, one-way ANOVA was performed. These methods were chosen based on the distribution of the data and the number of comparisons required. The level of significance was set at p < 0.05 for all tests.

### Creating figures and charts

Figures and charts were created using GraphPad Prism 10.2.2, Adobe Illustrator, Microsoft PowerPoint and BioRender. Each software tool was selected for its specific capabilities in graphical representation and design to ensure high-quality visual data presentation. Images were not altered except to enhance brightness, contrast, or introduce psuedo-color for fluorescence.

---

**Supplemental Figure 1.**
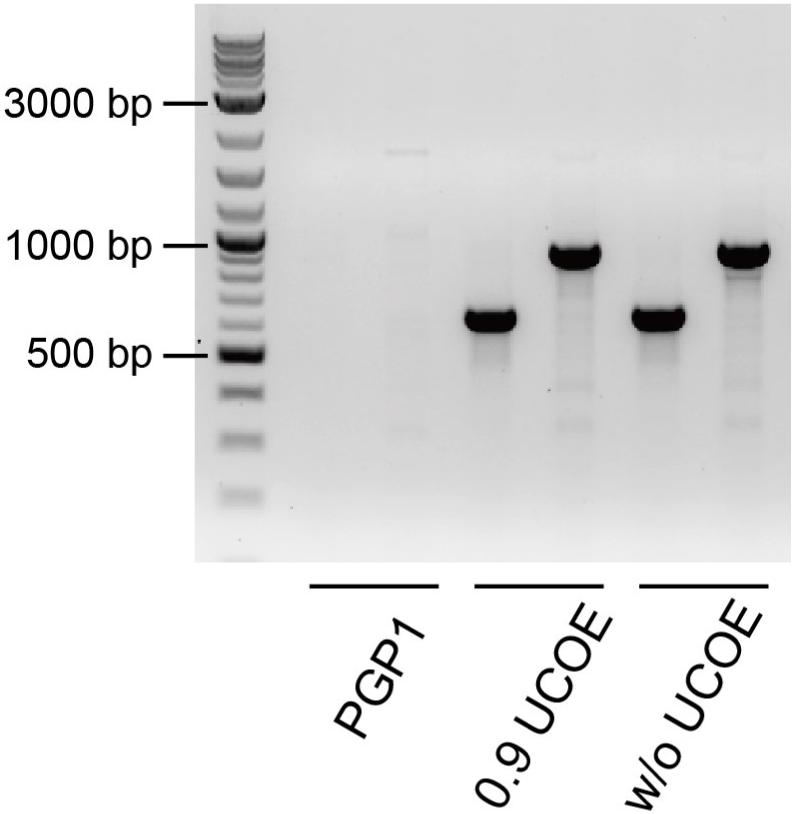
Junction PCR analysis of bulk cells with integrated gene circuits. Integration of the 0.9 UCOE-FOXN1 gene circuit (0.9 UCOE) and the without UCOE-FOXN1 gene circuit (w/o UCOE) into the Rogi1 target region in bulk cell populations was confirmed using PCR. PCR amplification revealed 650 bp and 921 bp bands corresponding to the 5’ and 3’ integration junctions, respectively.

**Supplemental Figure 2.**
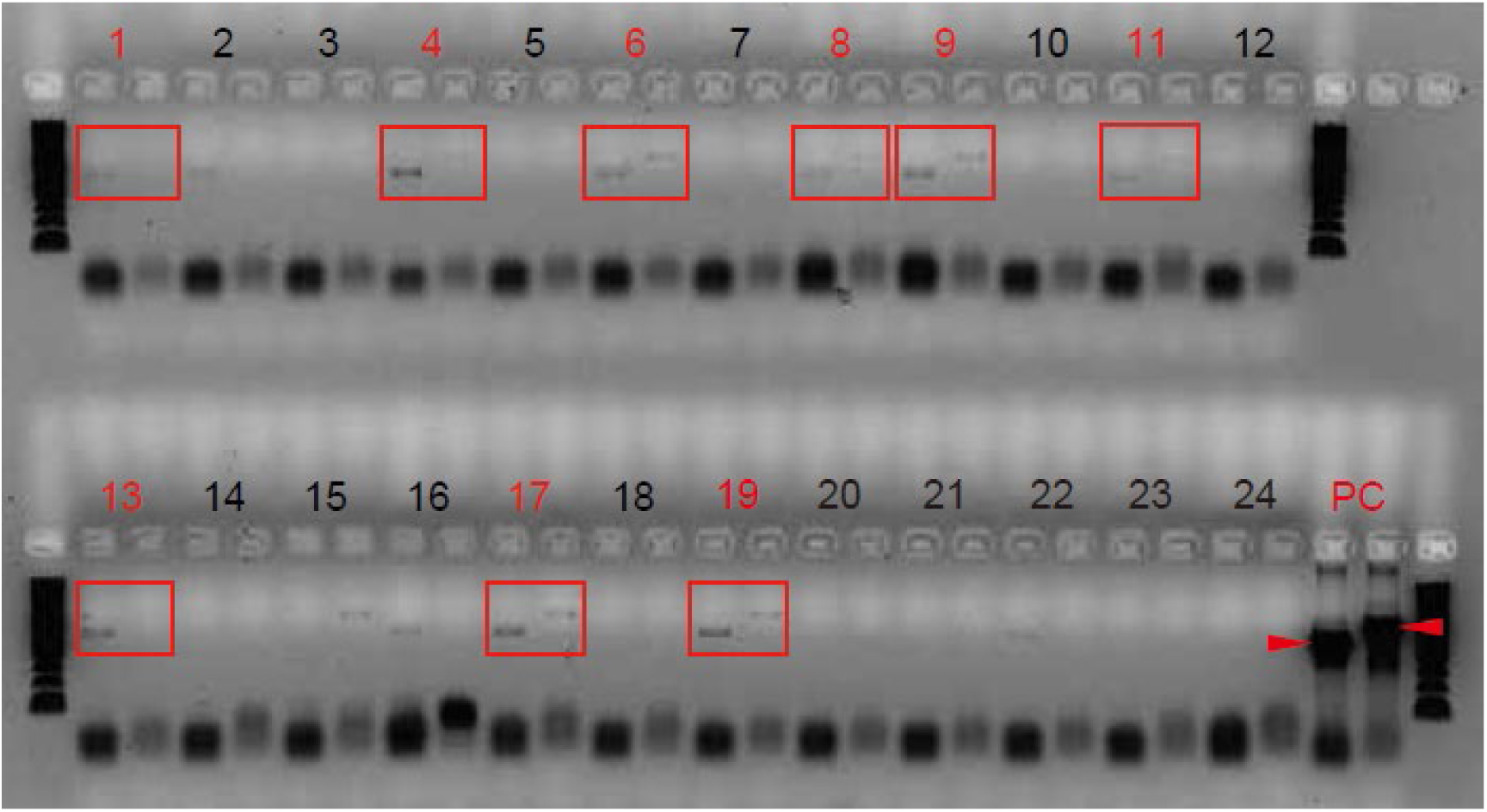
Junction PCR of iPS cells with integrated 0.9 UCOE-*FOXN1* gene circuit. A total of 24 colonies were pocked, among which 9 colonies (indicated in red) showed the correct PCR band. Abbreviations: PC, positive control.

**Supplemental Figure 3.**
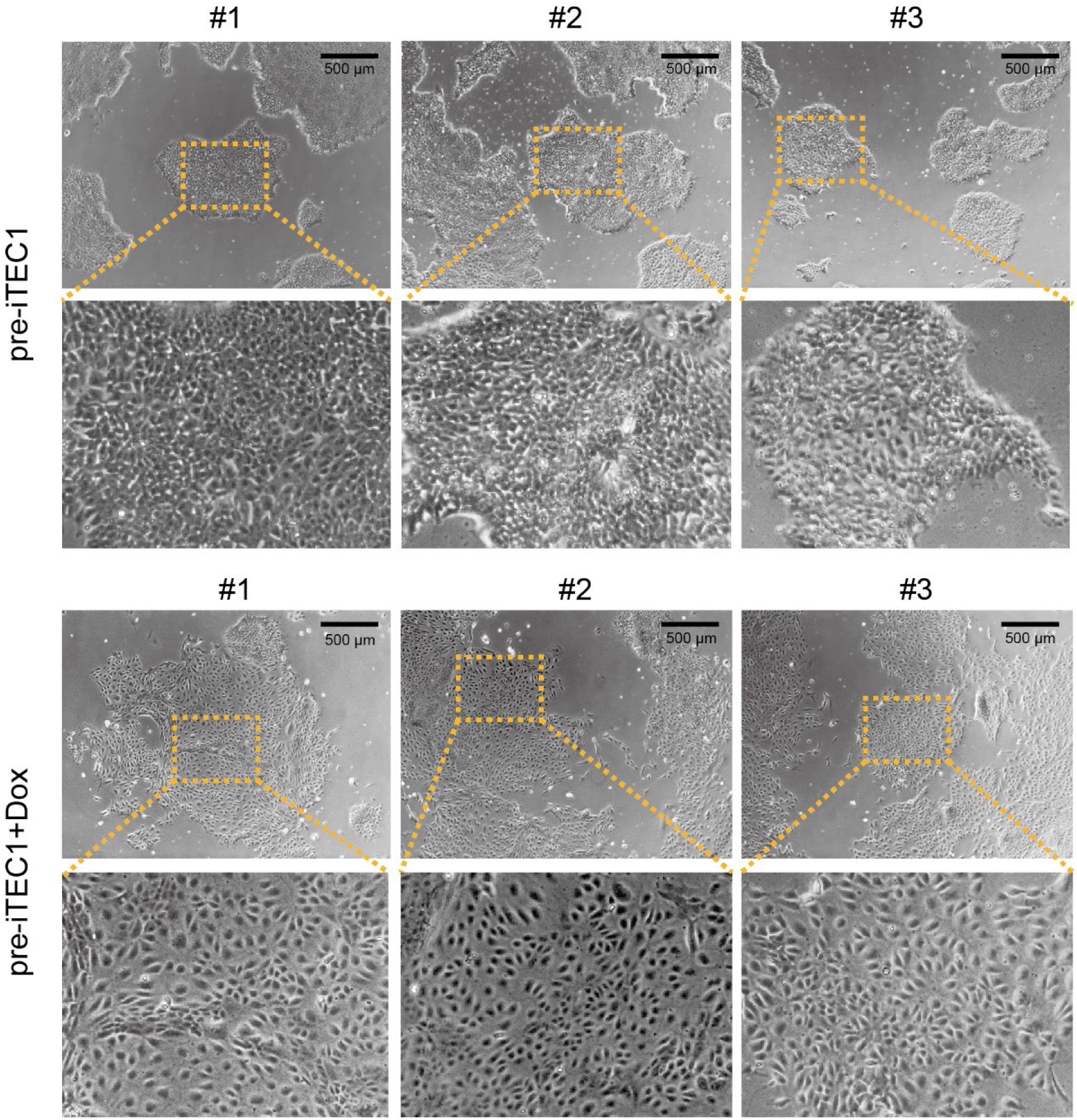
Morphological changes of pre-induced thymic epithelial cells (pre-iTEC) upon doxycycline treatment. Representative images showing morphological differences between untreated and doxycycline (Dox) -treated pre-iTEC cells. The untreated group (top two row) maintained a dense colony-like appearance characteristic of iPSCs, whereas the doxycycline-treated group (bottom two row) exhibited a looser and elongated form.

**Supplemental Figure 4.**
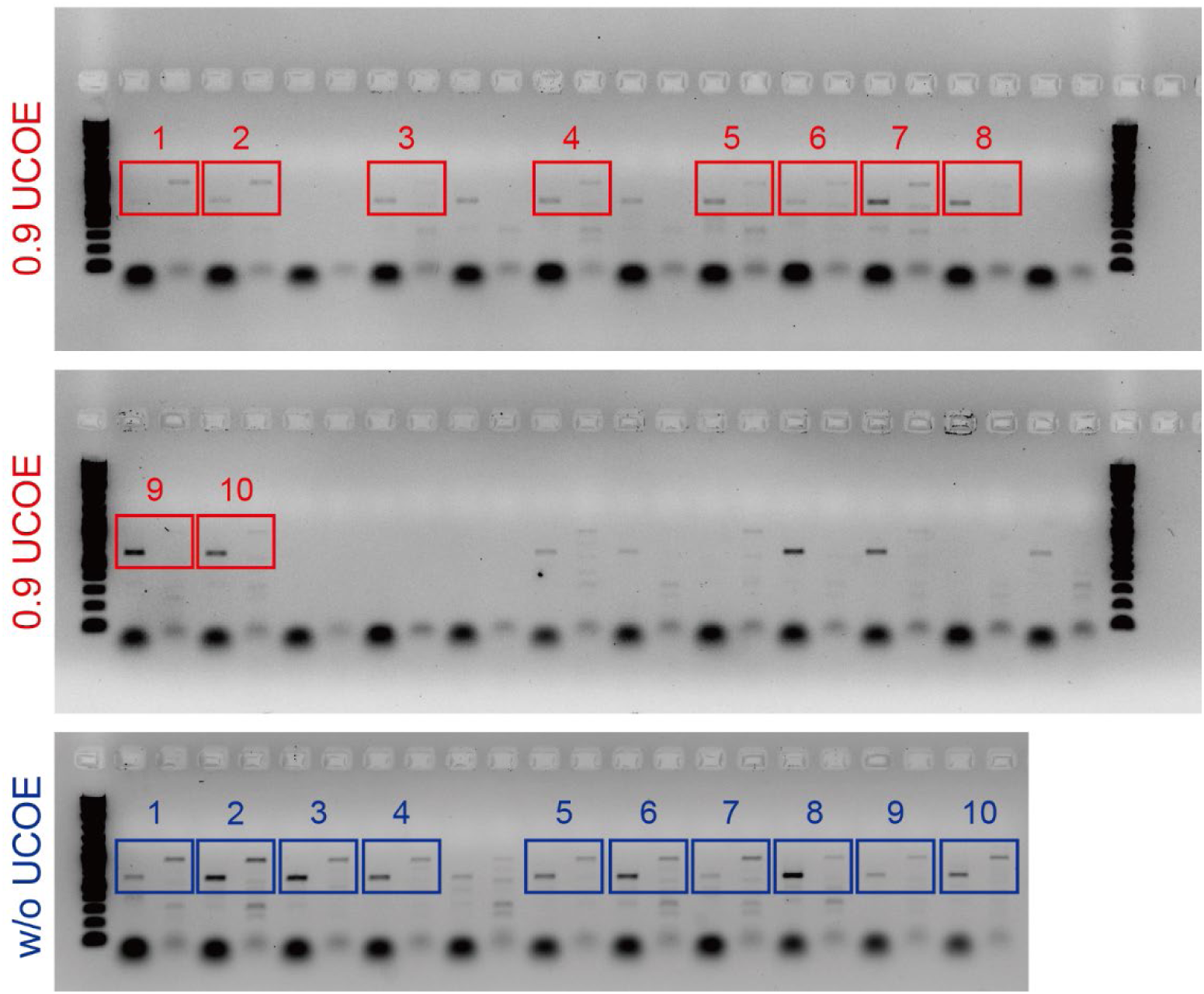
Junction PCR of single-cell-derived clones with integrated 0.9 UCOE-*FOXN1* or without UCOE-*FOXN1* circuits. Ten colonies each were picked from the 0.9 UCOE-*FOXN1* (0.9 UCOE) and without UCOE-*FOXN1* (w/o UCOE) groups. PCR was performed to confirm precise gene integration at the Rogi1 target site.

**Supplemental Figure 5.**
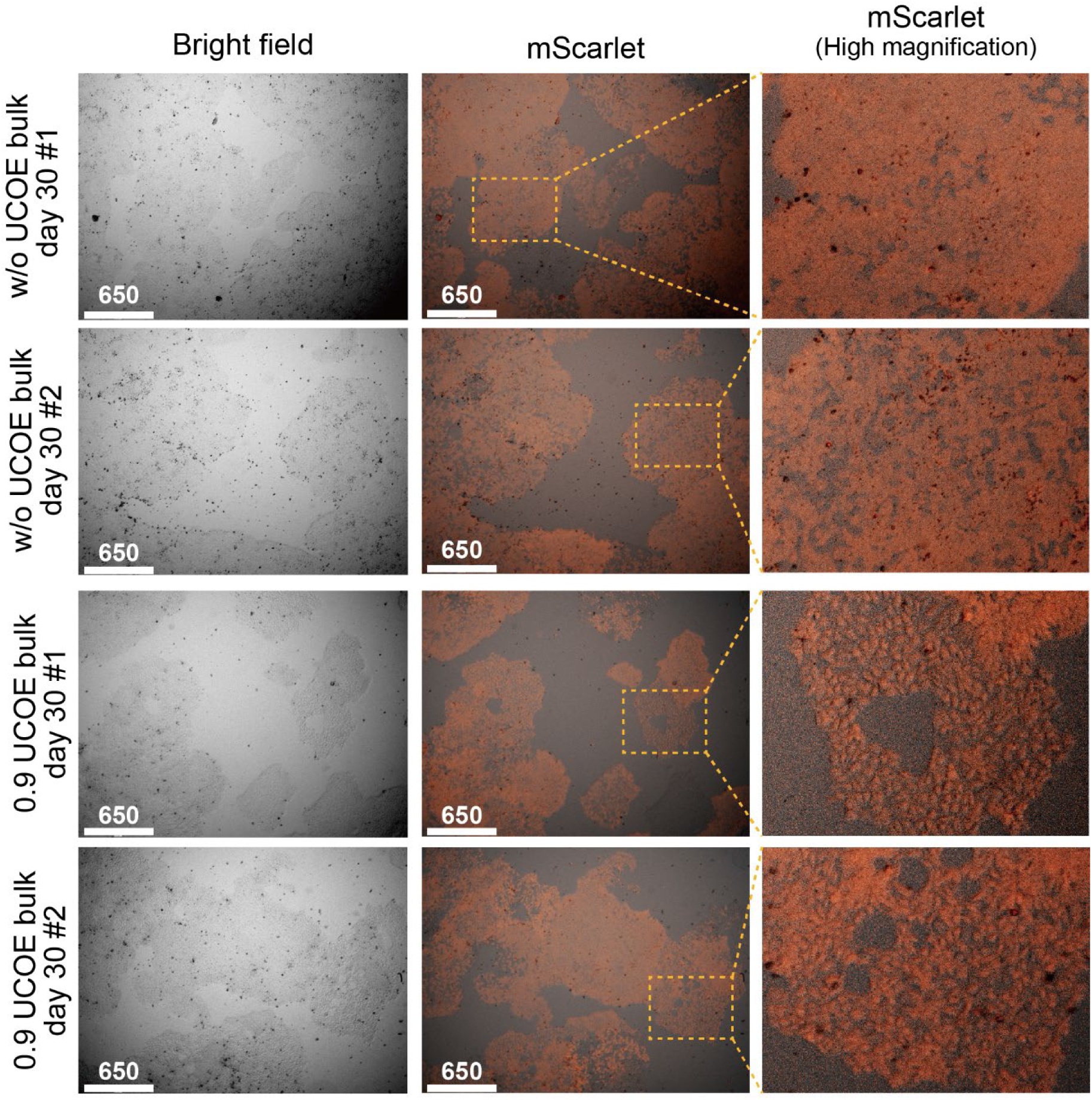
Morphological evaluation of bulk cell populations with integrated 0.9 UCOE-*FOXN1* (0.9 UCOE) or without UCOE-*FOXN1* (w/o UCOE) gene circuits after 30 days of culture. Representative images showing bulk cell populations cultured for 30 days in the presence of blasticidin but without doxycycline. The w/o UCOE-*FOXN1* group (top two rows) maintained a dense colony-like appearance, characteristic of iPS cells, even after 30 days. In contrast, the 0.9 UCOE-*FOXN1* group (bottom two rows) exhibited a looser and elongated form.

**Supplemental Figure 6.**
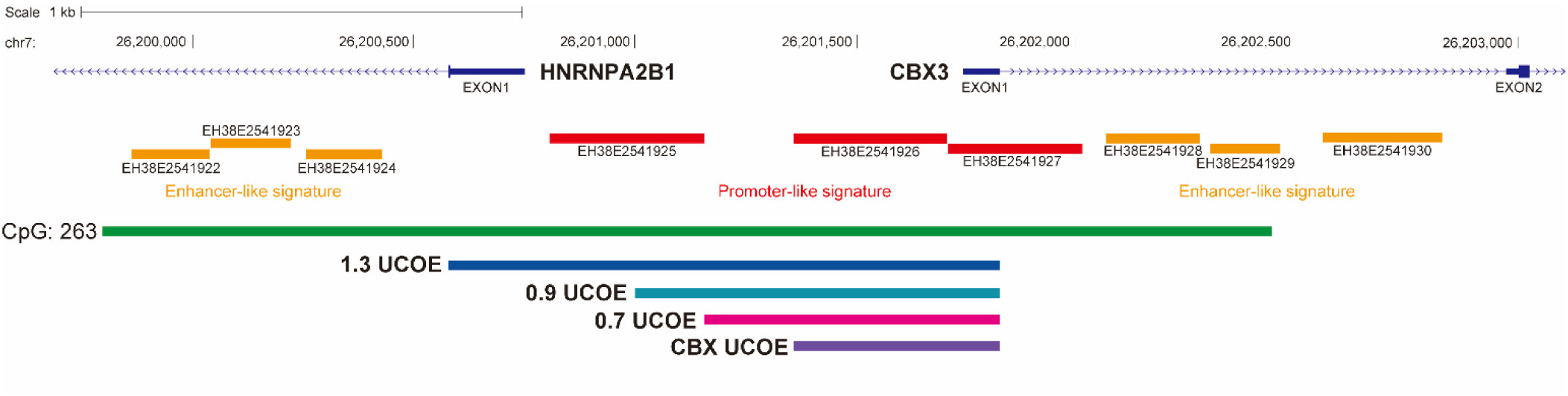
Design of UCOE fragments and their relationship to the *HNRNPA2B1* and *CBX3* genes, promoter regions, and enhancer elements. This figure illustrates the genomic structure of the *HNRNPA2B1* and *CBX3* genes in the human genome (GRCh38), using the UCSC Genome Browser to depict the surrounding regulatory elements. The red boxes indicate promoter regions, while the yellow boxes represent enhancer regions, and the green box marks the location of a CpG island. The 1.3 UCOE fragment spans from exon 1 of the HNRNPA2B1 gene to exon 1 of the *CBX3* gene, encompassing a promoter-like signature region between the two genes. The 0.9 UCOE fragment includes part of the upstream promoter region known as EH38E2541925 from *HNRNPA2B1*, extending to exon 1 of *CBX3*. The 0.7 UCOE fragment covers the region between EH38E2541925 and EH38E2541926, extending to the enhancer EH38E2541926 and exon 1 of *CBX3*. The smallest fragment, referred to as CBX UCOE, contains only the EH38E2541926 enhancer and exon 1 of *CBX3*.

**Supplemental Figure 7.**
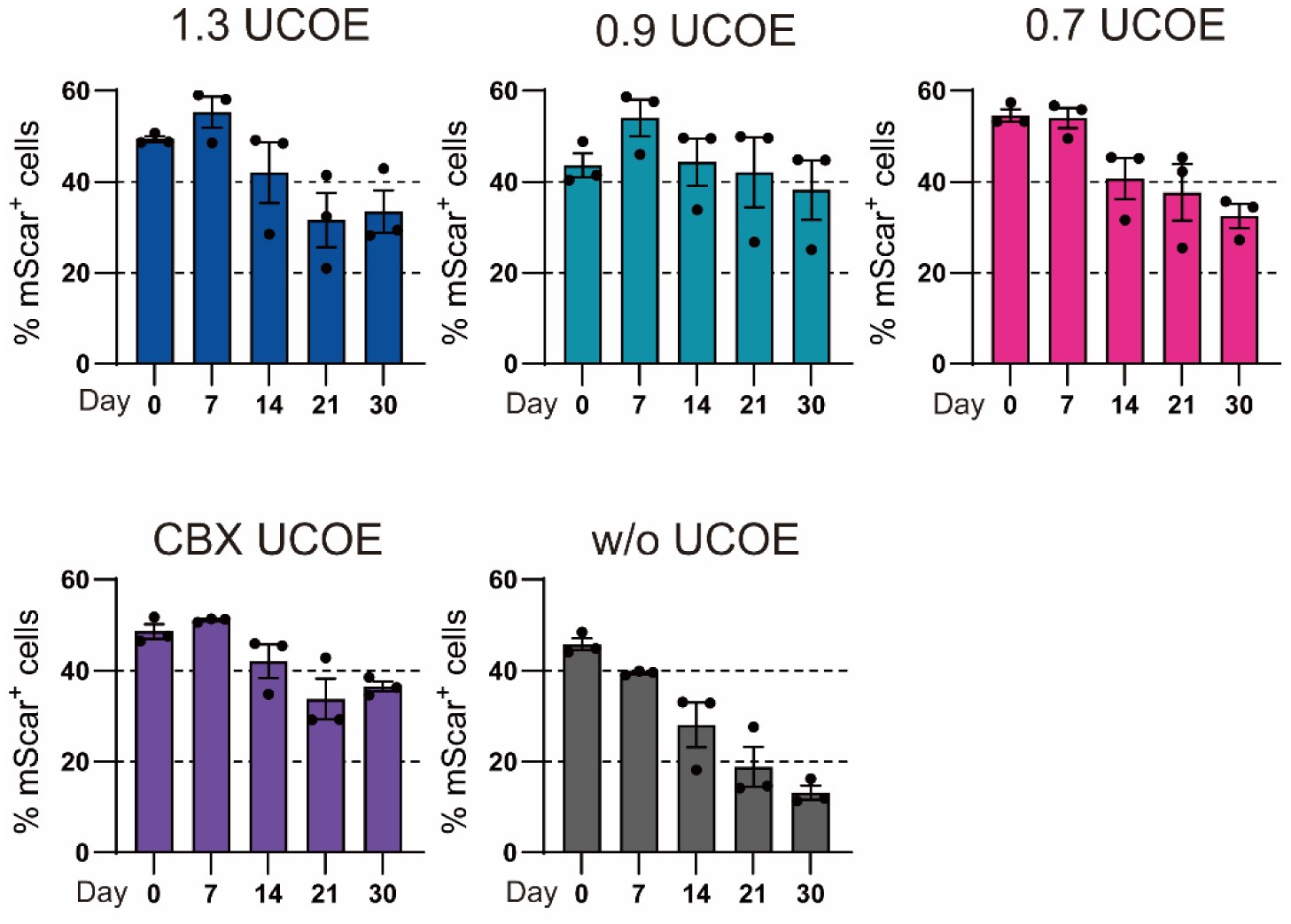
Flow cytometry analysis of RFP-positive cells in various UCOE fragment groups over a 30-day period without blasticidin. The RFP-positive (mScarlet+) cell percentage was evaluated in each UCOE fragment group on Days 0, 7, 14, 21, and 30 (n=3).

**Supplemental Figure 8.**
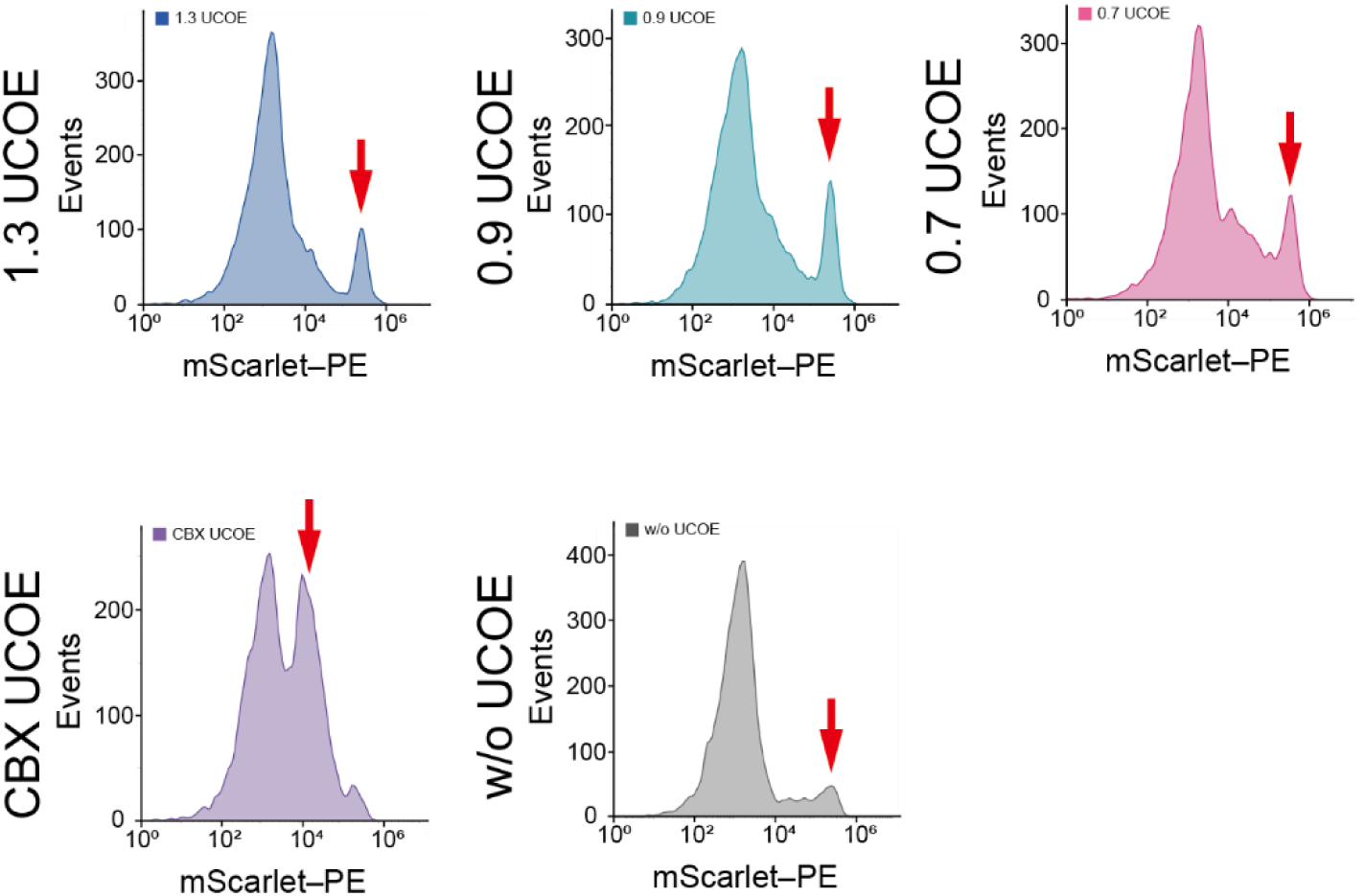
Differences in fluorescence intensity across various UCOE fragments. The fluorescence peak for each UCOE fragment group is indicated by a red arrow.

**Supplemental Figure 9.**
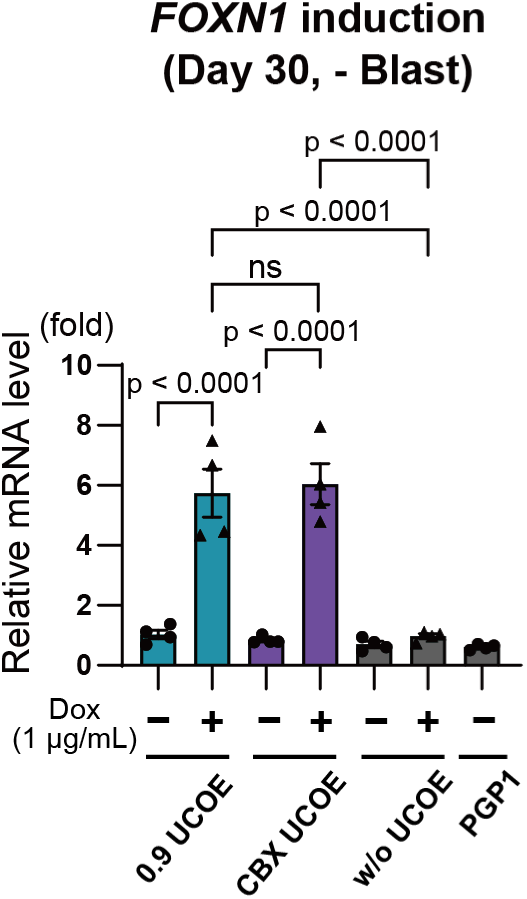
Evaluation of *FOXN1* transcription induction upon doxycycline treatment on Day 30. RT-qPCR analysis was used to assess *FOXN1* transcription levels in the 0.9 UCOE, CBX UCOE, and w/o UCOE groups after 30 days of culture in the absence of blasticidin, followed by doxycycline treatment (n=4). P values were calculated using one-way analysis of variance with Tukey’s honestly significant difference test. The data are presented as mean ± SEM. For detailed data, statistical analyses, and exact p-values, see source data file.

**Supplemental Figure 10.**
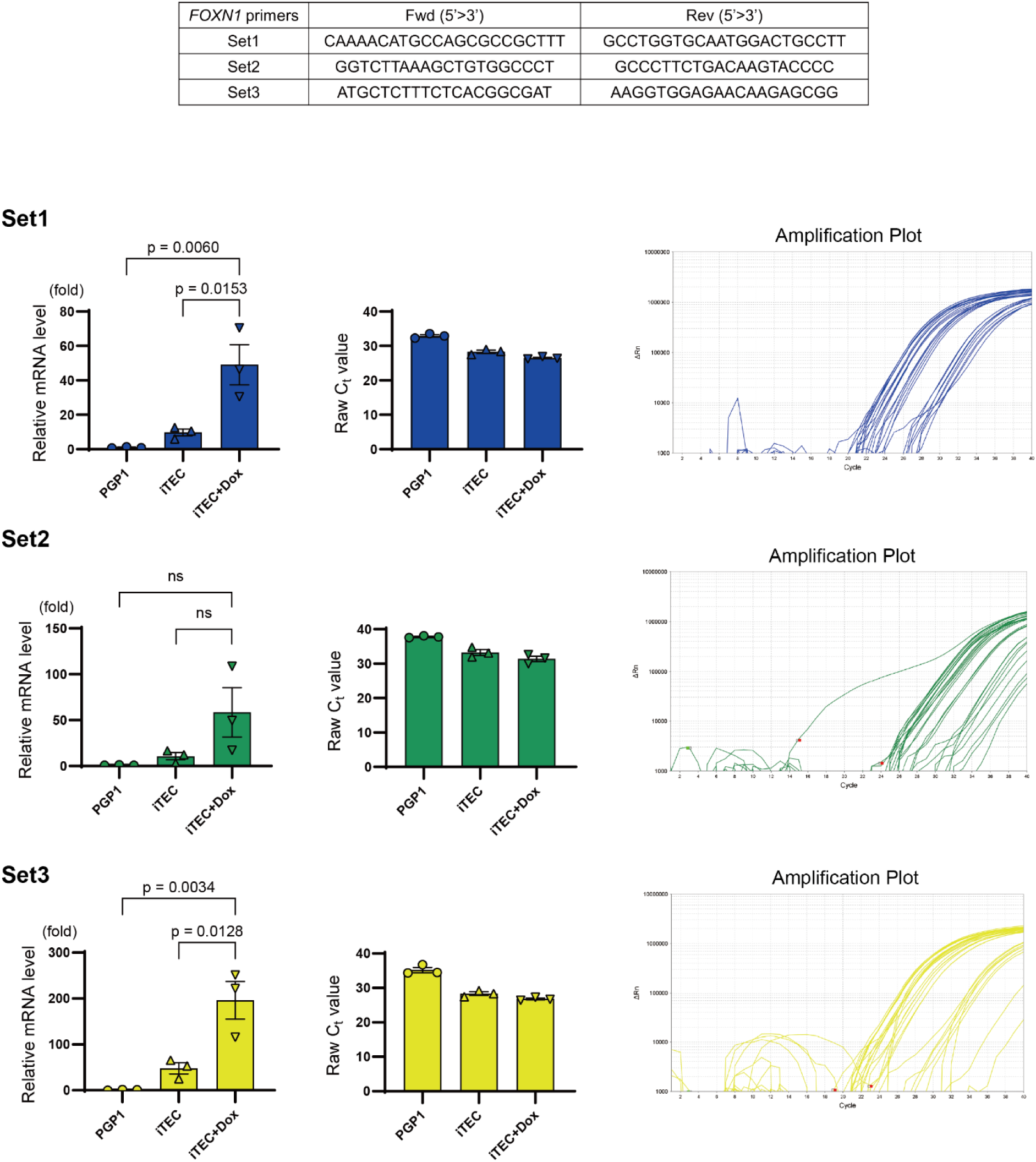
Evaluation of primer sets used for *FOXN1* leakage analysis. Three primer sets were designed for RT-qPCR targeting *FOXN1*. Relative mRNA levels normalized to the housekeeping gene *RPS29*, raw Ct values, and amplification plots are shown for each primer set.

**Supplemental Table 1.**
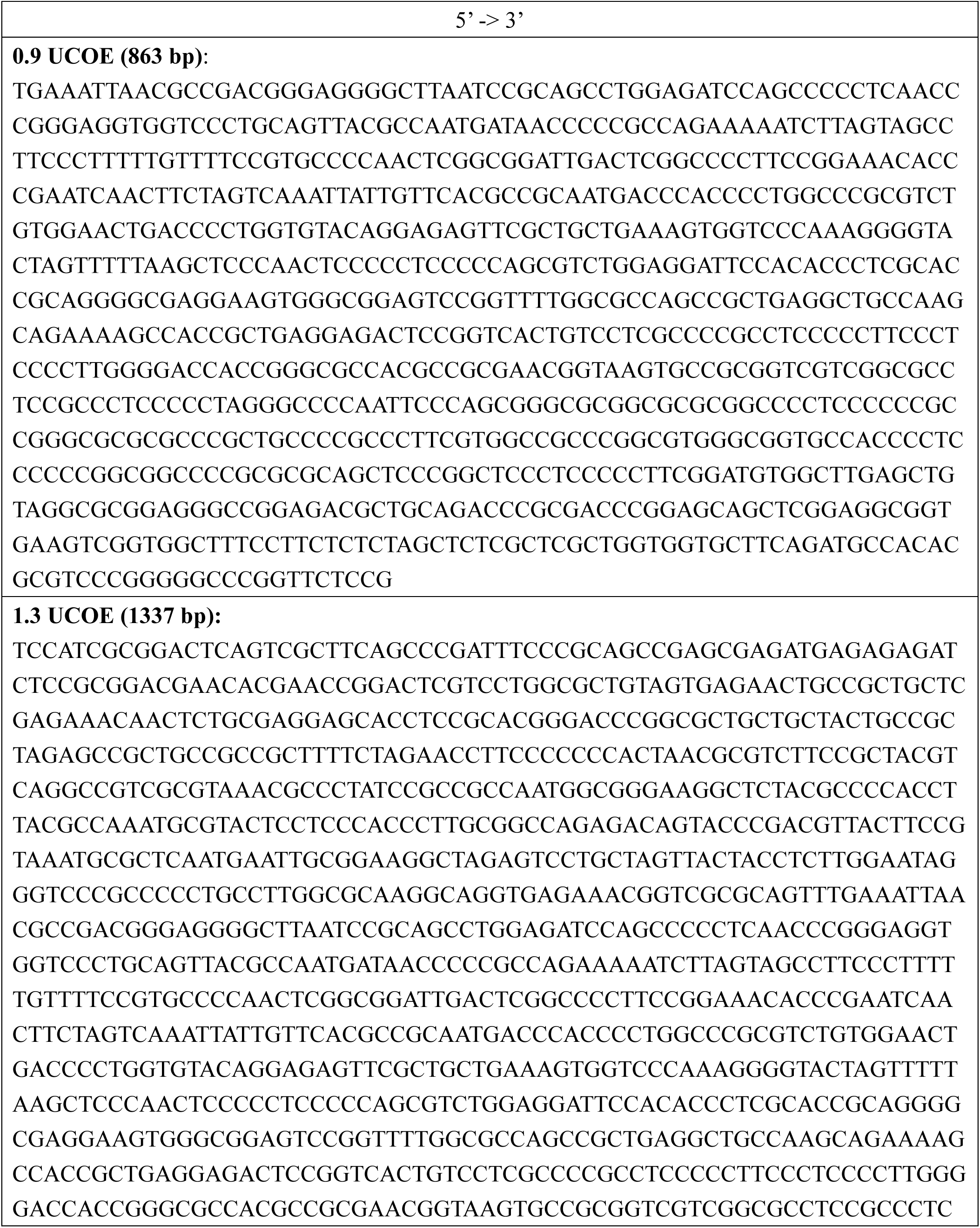
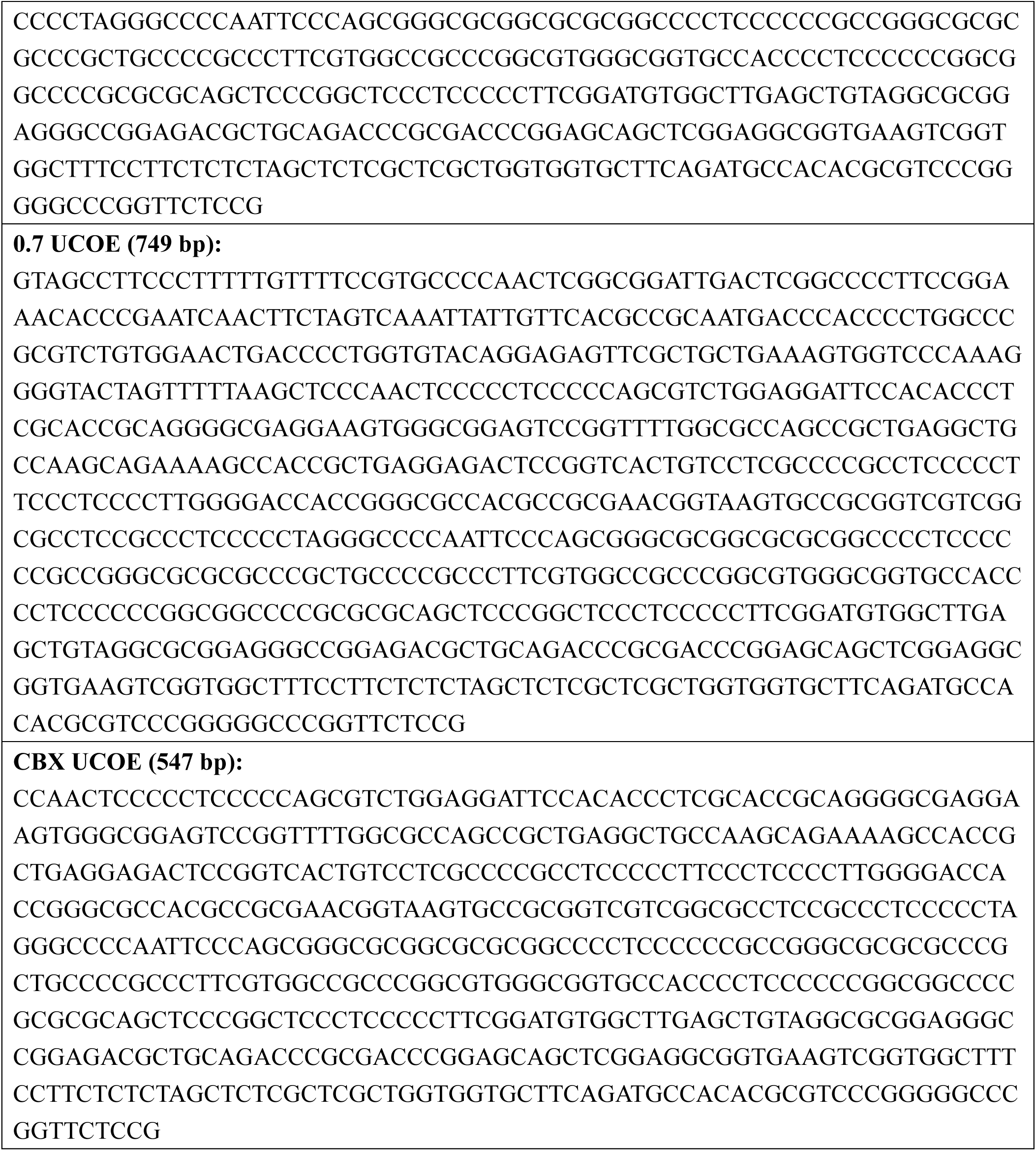
Sequence Information for Various A2UCOE Fragments. This table provides the sequence information for different A2UCOE fragments, including 0.9 UCOE, 1.3 UCOE, 0.7 UCOE, and CBX UCOE.

**Supplemental Table 2.**
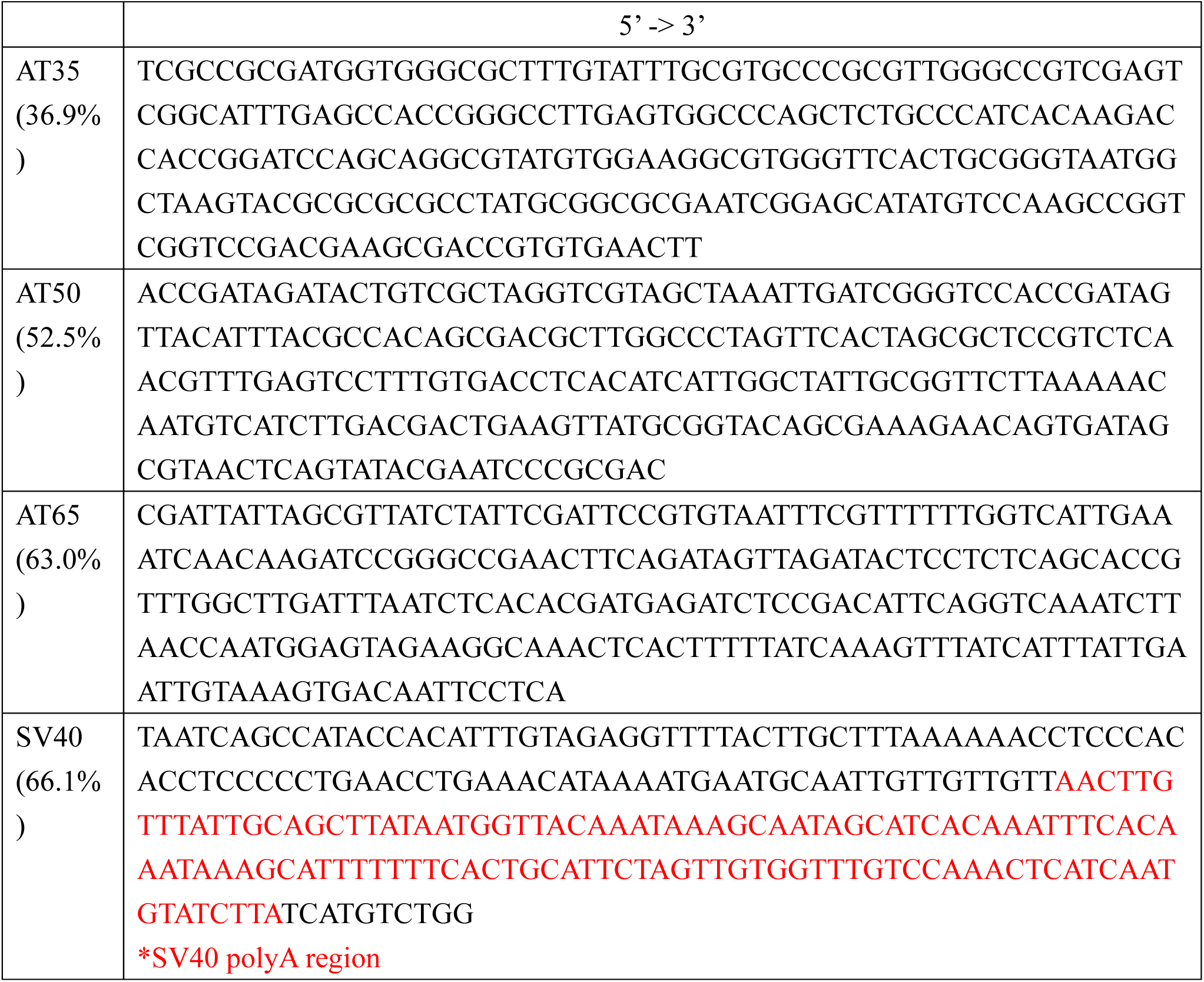
Sequence information for random AT-rich spacer sequences. This table provides the sequence information for the AT35, AT50, AT65, and SV40 spacer sequences. The SV40 polyadenylation (poly A) region is highlighted in red.

**Supplemental Table 3.**
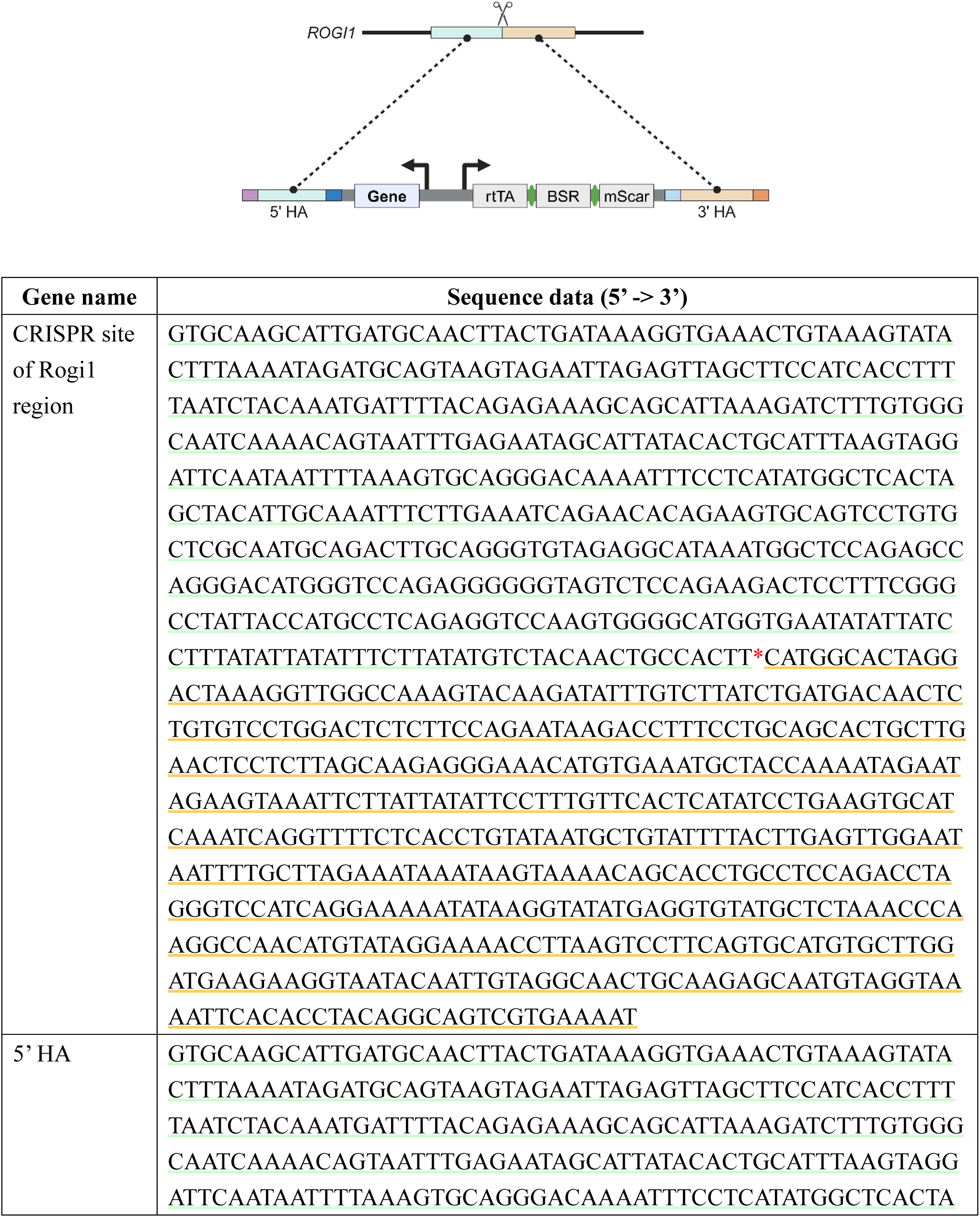
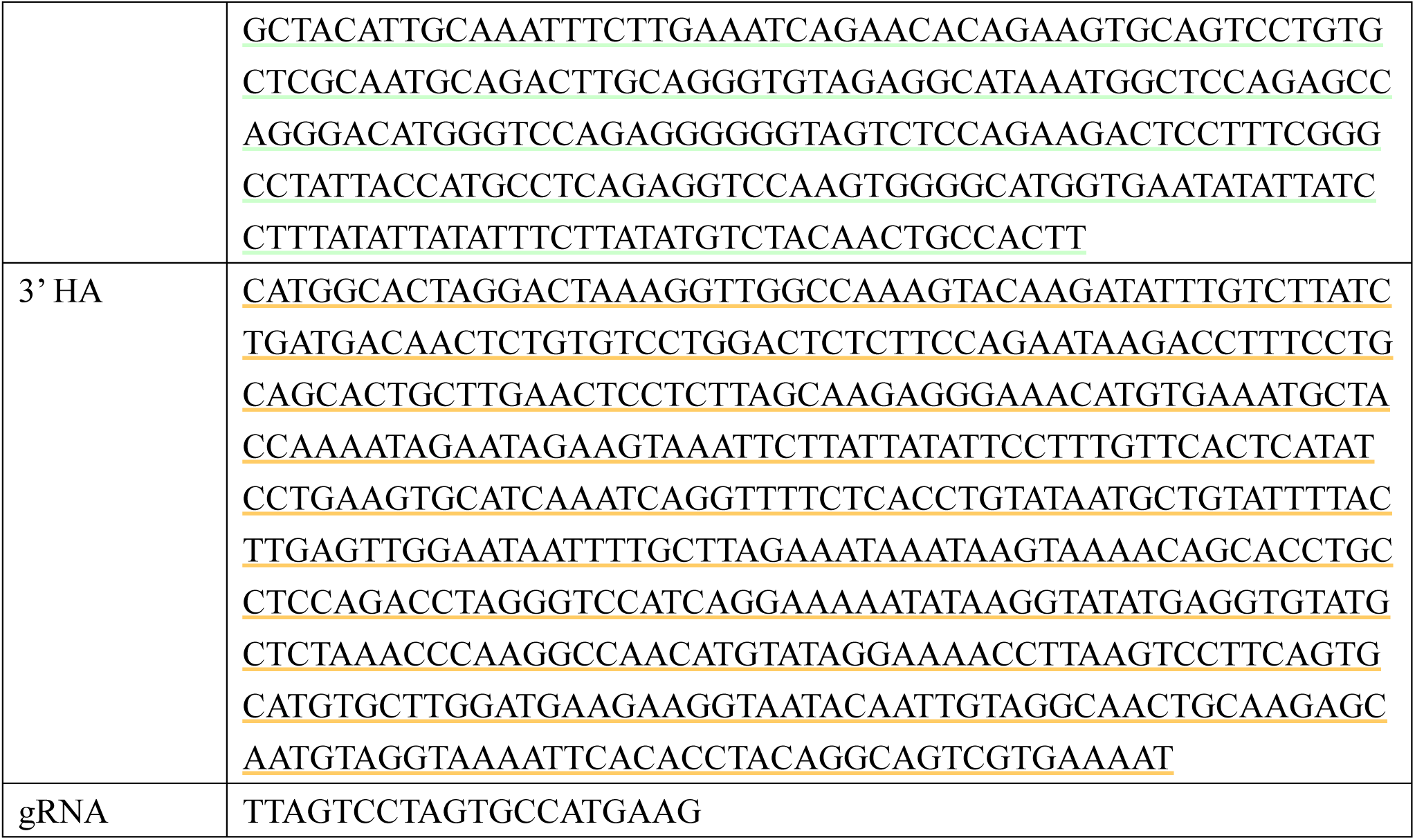
Sequence information for homology arms and gRNA for genomic integration at the Rogi1 site. This table provides the sequence information for the homology arms and gRNA used for CRISPR/Cas9-mediated cleavage at the Rogi1 site. The cleavage site is marked with an asterisk (*). The 5’ homology arm region is indicated by a green underline, and the 3’ homology arm region by a yellow underline.

**Supplemental Table 4.**
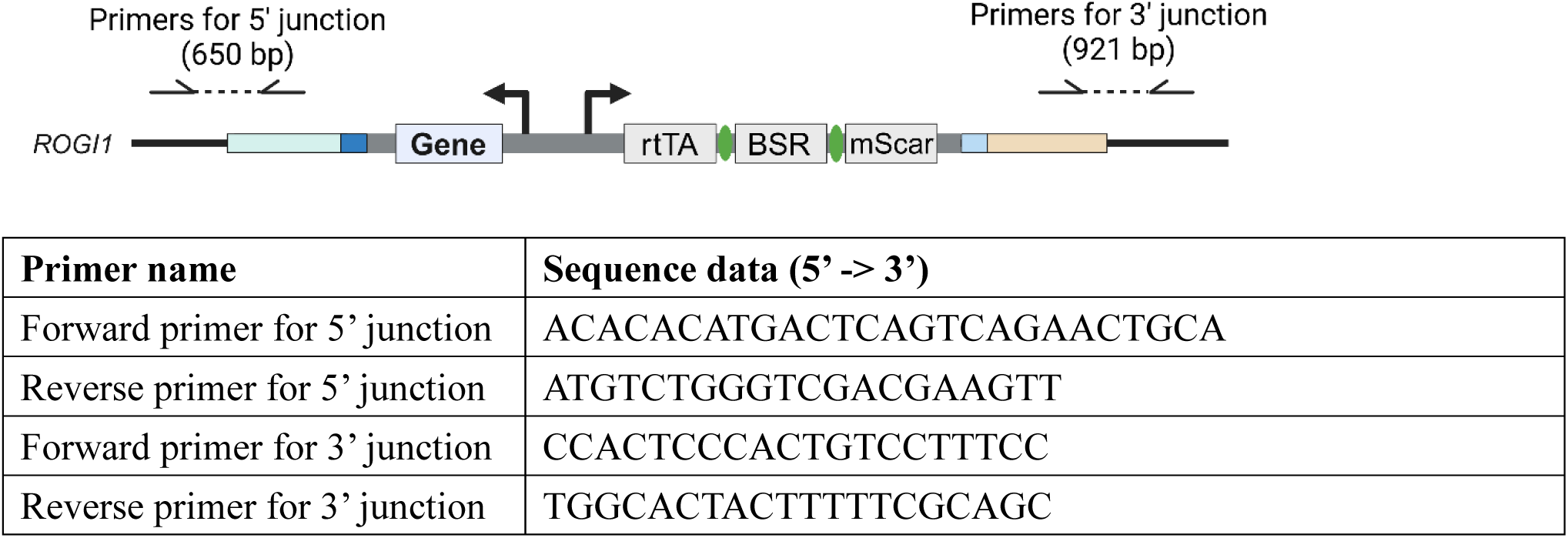
Primer sequences for junction PCR following genomic integration. This table lists the primer sequences used for junction PCR at the 5’ and 3’ junctions following genomic integration.

**Supplemental Table 5.**
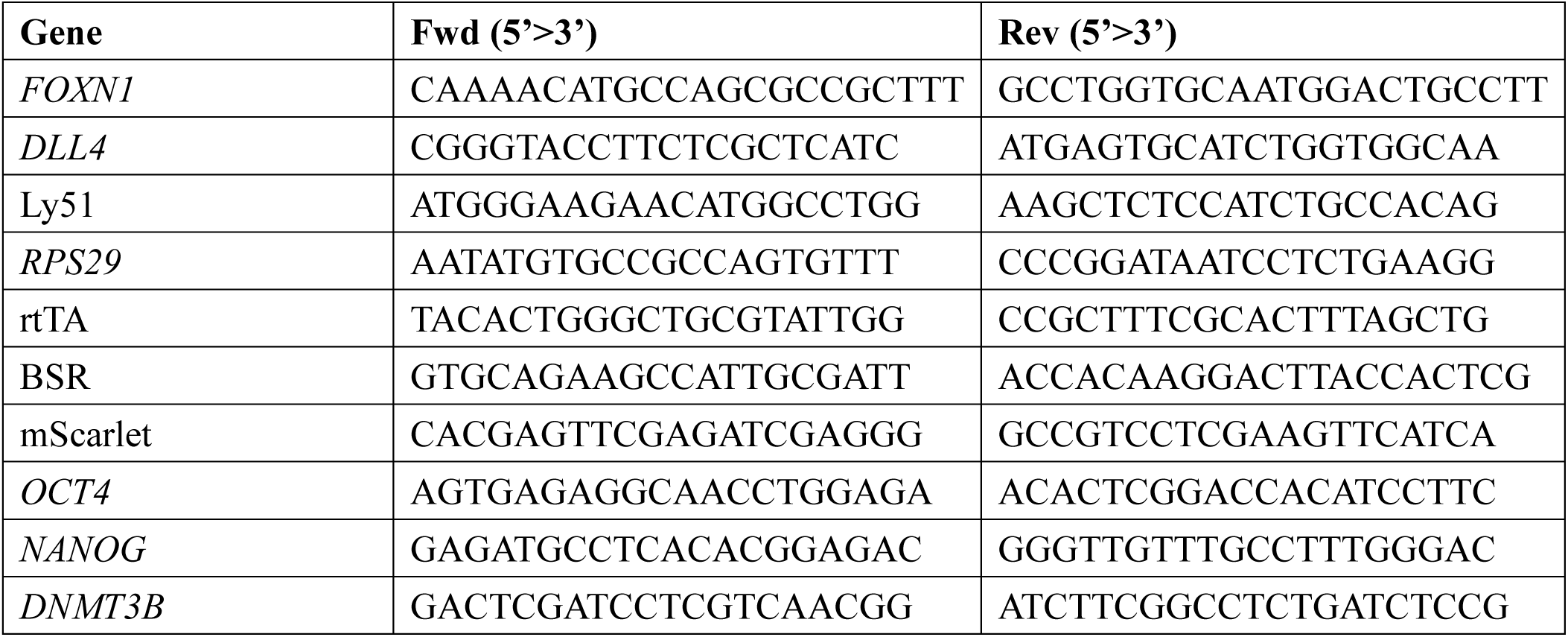
Primer sequences used for qPCR analysis. This table provides the sequences of primers used in qPCR experimen

## Notes

### Competing Interest Statement

The authors have declared no competing interest.

